# Meiosis-specific functions of kinetochore protein SPC105R required for chromosome segregation in *Drosophila* oocytes

**DOI:** 10.1101/2024.03.14.585003

**Authors:** Jay N. Joshi, Neha Changela, Lia Mahal, Tyler Defosse, Janet Jang, Lin-Ing Wang, Arunika Das, Joanatta G. Shapiro, Kim McKim

## Abstract

The reductional division of meiosis I requires the separation of chromosome pairs towards opposite poles. We have previously implicated the outer kinetochore protein SPC105R/KNL1 in driving meiosis I chromosome segregation through lateral attachments to microtubules and co-orientation of sister centromeres. To identify the domains of SPC105R that are critical for meiotic chromosome segregation, an RNAi-resistant gene expression system was developed. We found that SPC105R’s C-terminal domain (aa 1284-1960) is necessary and sufficient for recruiting NDC80 to the kinetochore and building the outer kinetochore. Furthermore, the C-terminal domain recruits BUBR1, which in turn recruits the cohesion protection proteins MEI-S332 and PP2A. Of the remaining 1283 amino acids, we found the first 473 are most important for meiosis. The first 123 amino acids of the N-terminal half of SPC105R contain the conserved SLRK and RISF motifs that are targets of PP1 and Aurora B kinase and are most important for regulating the stability of microtubule attachments and maintaining metaphase I arrest. The region between amino acids 124 and 473 are required for two activities that are critical for accurate chromosome segregation in meiosis I, lateral microtubule attachments and bi-orientation of homologs.

**Significance Statement:**

- Kinetochore proteins regulate meiosis specific functions. SPC105R is a central regulator of kinetochore function but its role in meiosis is not well understood.
- We identified regions of SPC105R that regulate key meiosis I functions, including fusing sister centromeres and the way the kinetochore interacts with the microtubules.
- SPC105R is a hub that recruits several proteins to regulate kinetochore activity. Future work will involve identifying the proteins recruited by SPC105R that mediate these functions in meiosis.

## Introduction

Kinetochores are large protein complexes built on top of the centromeres that interact with microtubules and regulate cell cycle progression (McAinsh and Marston, 2022; Musacchio and Desai, 2017). The conserved KMN network is required for kinetochore-microtubule attachments *in vivo* and is composed of three groups of proteins: Spc105/KNL1, the Mis12 complex, and the Ndc80 complex (Przewloka and Glover, 2009). SPC105R is the *Drosophila* homolog of KNL1 (Schittenhelm et al., 2009) and is required for outer kinetochore assembly in oocytes (Radford et al., 2015). Within the KMN network, two microtubule binding sites have been identified, in Ndc80 and Spc105/KNL1 (Cheeseman et al., 2006). In addition to facilitating microtubule attachments, KNL1 serves as a scaffold to recruit spindle assembly checkpoint proteins (Caldas and DeLuca, 2014). Being involved in all these activities, Spc105R/KNL1 has a critical role in regulating cell division.

In meiosis I, homologous chromosomes segregate into daughter cells, resulting in a reduction of ploidy, rather than segregation of sister chromatids during meiosis II or mitosis. Thus, homologous chromosome segregation in meiosis I involves unique chromosome-based mechanisms (Paliulis and Nicklas, 2000). SPC105R regulates two unique meiosis 1 functions: co-orientation of sister chromatids and bi-orientation of homologous chromosomes (Radford et al., 2015). Co-orientation is the process of fusing sister kinetochores during prometaphase I. Regulation of cohesion loss is a critical component that differentiates the two divisions. Arm cohesion is lost in meiosis I, but cohesion around the centromeres must be maintained until meiosis II (Ogushi et al., 2021). Failure to fuse sister kinetochores results in merotelic attachments, with one pair of sister chromatids attached to both spindle poles (Nasmyth, 2015; Watanabe, 2012). We previously found that co-orientation depends on two independent mechanisms, regulation of end-on microtubule attachments and centromeric cohesion (Wang et al., 2019). How SPC105R regulates meiotic cohesion, and microtubule attachments, is not known.

Bi-orientation is a critical part of pro-metaphase I that establishes how homologous chromosome pairs segregate at anaphase I. Pairs of homologous chromosomes, joined by chiasmata, bi-orient on a bipolar meiotic spindle by having their kinetochores attach to microtubules from opposite poles (Hughes et al., 2018). There are, however, important biological differences between oocytes and other cell types, such as the absence of centrosomes (Dumont and Desai, 2012; Kitajima, 2018; Mihajlovic and FitzHarris, 2018; Radford et al., 2017). Understanding these differences and how they impact the mechanism of biorientation oocytes might help explain the high error rate in human oocytes (Webster and Schuh, 2017).

Our work has shown that bi-orientation on an acentrosomal spindle depends on two modes of kinetochore-microtubule attachments: lateral attachments that depend on SPC105R, and stable end-on attachments that depend on NDC80 (Radford et al., 2015). The dependence of oocytes on lateral attachments for bi-orientation is likely conserved (Dumont et al., 2010; Wignall and Villeneuve, 2009) and present in other cell types (Itoh et al., 2018a; Magidson et al., 2015). During prometaphase in mouse oocytes, the chromosomes congress to the central spindle, which enhances the rate of bi-orientation by bringing kinetochores into the vicinity of a high density of microtubule plus ends (Kitajima et al., 2011; Magidson et al., 2011). *Drosophila* oocytes have a robust central spindle that depends on the kinesin-6 Subito, is composed of several proteins including the Aurora B kinase (Costa and Ohkura, 2019; Jang et al., 2005), and is required for error correction and bi-orientation during pro-metaphase (Das et al., 2018; Jang et al., 2007). Regulating the transition from lateral to end-on attachment appears to be critical for avoiding bi-orientation defects in mammals (Yoshida et al., 2015) and *Drosophila* (Głuszek et al., 2015).

We initiated this study of SPC105R in oocytes to understand how the meiosis-specific functions of kinetochore co-orientation and bi-orientation are regulated. To do this, we generated an RNAi-resistant functional *Spc105R* transgene and used it as a platform to study mutants with deletions of specific domains. These results show that the C-terminal domain is sufficient for recruiting outer kinetochore proteins and establishing end-on attachments. It also recruits cohesion protection proteins, which are required to fuse the sister kinetochores during meiosis I. The rest of SPC105R (aa 1-1283) regulates end-on attachments and bi-orientation. We identified one region in particular, between 123 and 473, that is required for bi-orientation of homologous chromosomes. Our evidence suggests this region is required for lateral microtubule interactions and regulating the transition from lateral to end-on attachments.

## Results

### Using RNAi-resistant transgenes to study SPC105R function

There are two *Spc105R* isoforms predicted from the genome sequence: *Spc105R^A^* and *Spc105R^B^* (Figure S 1A). Alternative splicing results in different translation start sites, such that the sequence MNANKRRSSLRK in the B form is replaced with MVDLLFLQLRK in the A form. The only known full length cDNA encodes the A form, but multiple 5’ cDNA sequences from RACE experiments correspond to the B form (Gramates et al., 2022). Furthermore, RT-PCR amplification specific to mRNA for each isoform from wild-type female ovaries confirmed the presence of both A and B isoforms (Figure S 1B). Both isoforms could be detected at the kinetochores when expressed using MYC-tagged transgenes (Figure S 1C). These results suggest that both isoforms are incorporated into the kinetochore.

In order to study the function of *Spc105R* in meiosis, we constructed RNAi-resistant transgenes that contained silent mutations to the bases targeted by the shRNA *GL00392* (Figure 1A). While Drosophila females expressing *GL00392* in the ovary using *matα* are sterile, females expressing the RNAi-resistant B-form of SPC105R, *Spc105R^B^,* and *GL00392* (to be referred to as *Spc105R^B^, Spc105R^RNAi^* oocytes) are fertile (Table 1). *Spc105R^B^, Spc105R^RNAi^* females were more fertile than *Spc105R^A^, Spc105R^RNAi^* females, suggesting that SPC105R^B^ is the more important isoform for providing functional rescue of Spc105R. *Spc105R^B^*rescued the lethality caused by ubiquitous expression of *GL00392* using *P{w[+mC]=tubP-GAL4}LL7* (referred to as *Tub:GAL4*) (Figure 1B). This lethality was also suppressed by expression of *Spc105R^A^*, consistent with the conclusion that both isoforms are functional in mitosis.

**Figure 1:**
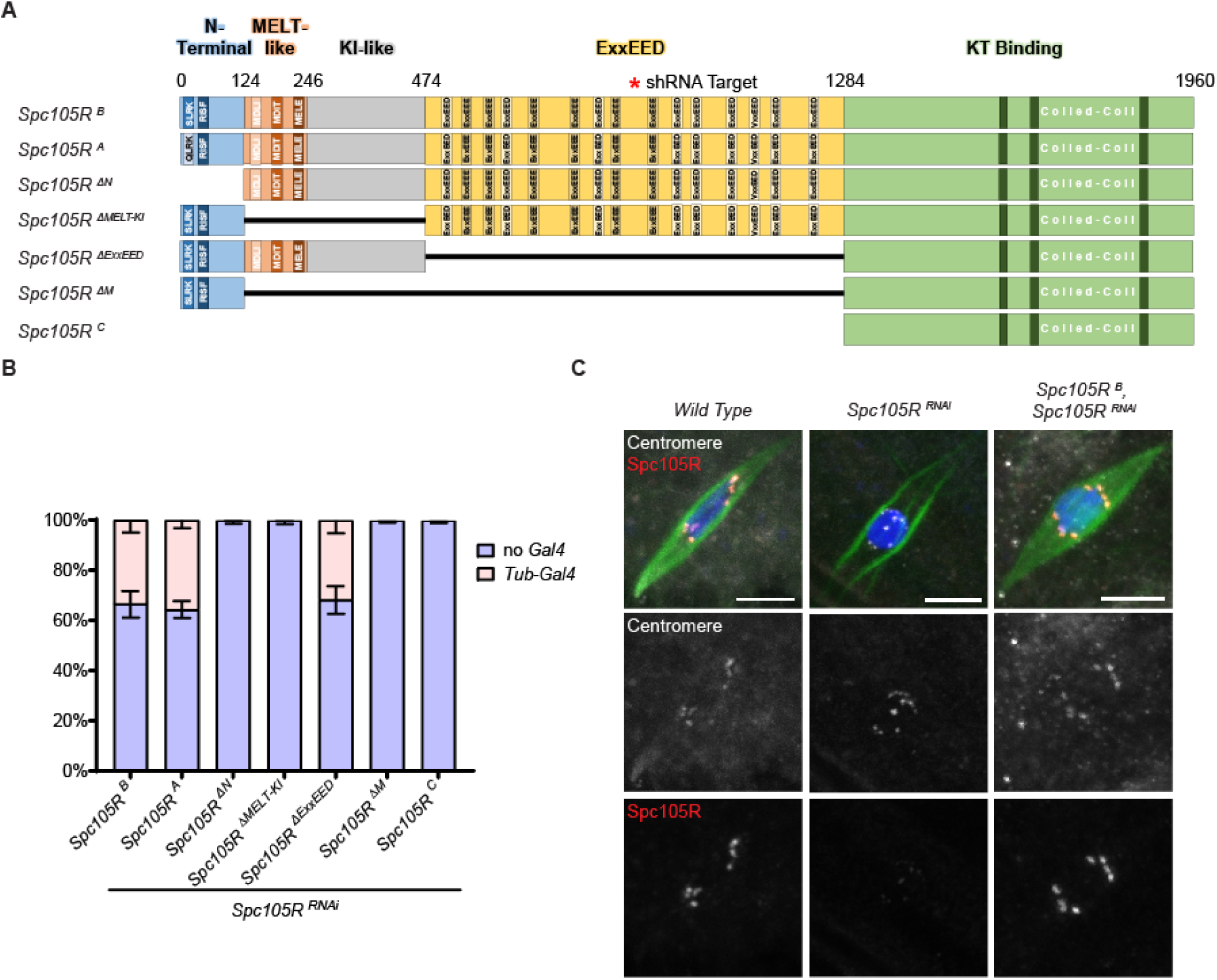
Spc105R^B^ can rescue most wild-type SPC105R functions. (A) A schematic of the two known *Drosophila Spc105R* isoforms and mutants used in this study. The first 9 amino acids in SPC105R^B^ are changed from MNANKRRSS to MVDLLFLQ in SPC105R ^A^. The coordinates on the schematic represent the first amino acid of each domain. The N-terminal includes the SLRK and RISF motifs and is where the two isoforms differ. Following this is a domain with three MELT-like motifs, a region that contains two KI-like repeats, a central domain containing repeats with the consensus ExxEED, and the C-terminal region containing coiled-coil motifs. All transgenes included missense mutations to make them resistant to the shRNA *GL00392*. (B) Viability of *Spc105R* mutants, all in a *Spc105R^RNAi^* background. Data shows the relative amounts of progeny expressing the *Spc105R^RNAi^*and mutants (*Tub-Gal4)* to siblings that did not (no *Gal4*) (n> 200). (C) Confocal images of w*ild-type*, *Spc105R^RNAi^*, and *Spc105R ^B^*, *Spc105R^RNAi^* oocytes. Merged images show DNA (blue), Tubulin (green), centromeres (white), and SPC105R (red). Centromeres and SPC105R are shown in separate channels. Scale bars are 5 μm.

**Table 1:** Fertility and nondisjunction in *Spc105R* domain mutant females.

| *Spc105R<sup>RNAi</sup>* + | NDJ (%) | Total |
| --- | --- | --- |
| *Spc105R<sup>B</sup>* | 0.1 | 2347 |
| *Spc105R<sup>A</sup>* | 0.0 | 59 |
| *Spc105R<sup>ΔN</sup>* | -- | sterile |
| *Spc105R<sup>ΔExxEED</sup>* | 10.3 **** | 604 |
| *Spc105R<sup>ΔM</sup>* | -- | sterile |
| *Spc105R<sup>ΔMELT-KI</sup>* | -- | sterile |
| *Spc105R<sup>C</sup>* | -- | sterile |
| No RNAi |  |  |
| *Spc105R<sup>B</sup>* | 0.5 | 2688 |
| *Spc105R<sup>A</sup>* | 0.3 | 1051 |
| *Spc105R<sup>ΔN</sup>* | -- | sterile |
| *Spc105R<sup>ΔExxEED</sup>* | 1.5 | 528 |
| *Spc105R<sup>ΔM</sup>* | -- | Sterile |
| *Spc105R<sup>ΔMELT-KI</sup>* | 5.6 **** | 460 |
| *Spc105R<sup>C</sup>* | -- | sterile |

Females expressing the indicated transgene were either in a *Spc105R<sup>RNAi</sup>* or wild-type
(dominant) background. Significant levels of bi-orientation defects ( $p < 0.001$ , Fishers
exact test) are shown by \*\*\*\*. NDJ = nondisjunction

Oocytes lacking SPC105R are characterized by the absence of several kinetochore proteins and microtubule attachments (Radford et al., 2015; Wang et al., 2019). The absence of kinetochore-microtubule attachments (KT-MT) results in a “hollow spindle” phenotype in which the centromeres lack an association with the microtubules (Figure 1C, Figure S 2A). All the microtubules are interpolar, with the plus-ends of microtubules from opposite poles making antiparallel overlaps in the center of the spindle. *Spc105R^B^*rescued the attachment defects present in *Spc105R^RNAi^* oocytes (Figure 1C). All the centromeres were observed to make contacts with microtubules, with the majority making end-on attachments to the plus ends of kinetochore microtubules. *Spc105R^B^* also rescued SPC105R localization to the kinetochore, which was absent in *Spc105R^RNAi^* oocytes (Figure 1C, Figure S 2A,B). These results show that *Spc105R^B^* can provide the meiotic and mitotic functions of SPC105R. Thus, throughout this analysis we used *Spc105R^B^* as our wild-type control.

Based on sequence comparisons and studies in mitotic embryo cells (Schittenhelm et al., 2009), we divided SPC105R into five regions (Figure 1A). The N-terminal domain includes potential Aurora B phosphorylation, PP1, and microtubule binding sites (Audett et al., 2022). Next to the N-terminal domain is a region containing three motifs that are related to the MELT consensus (MDLI, MDIT and MELE). This is followed by a disordered region that contains two sequences similar to the KI motifs of vertebrate orthologs (Audett et al., 2022; McGory et al., 2024). The C-terminal domain was previously shown to be sufficient for recruiting the NDC80 complex in mitotic cells (Schittenhelm et al., 2009).

The central region of the protein contains 15 repeats with the consensus ExxEED. This consensus is found in several *Drosophila* species (Tromer et al., 2015) although many organisms, including mammals, have repeats of the MELT motif in this domain. The *Drosophila* ExxEED repeats have been proposed to be phosphomimetic derivatives of the MELT motif. Consistent with this, there is a threonine 5 amino acids upstream of the ExxEED consensus in 14/15 of the repeats, and the 15^th^ is a serine. This threonine is a Aurora B phosphorylation site (Audett et al., 2022) and is part of the consensus KxRxTLL that is related to the TΩ motif upstream of the MELT motif in other organisms (Tromer et al., 2015).

Five deletion mutations, one for each domain, were made on the *Spc105R^B^*sequence. Each mutant protein was detected at the centromeres using anti-myc antibodies (Figure S 1). However, three mutants were detected by myc but not the anti-SPC10R antibody. The SPC105R antibody was raised against the first 400 amino acids of SPC105R (Schittenhelm et al., 2009).. Thus, it is expected that SPC105R^C-^would not be detected because that region was deleted. However, the absence of staining in *Spc105R^ΔM^ Spc105R^RNAi^*and *Spc105R^ΔMELT-KI^ Spc105R^RNAi^*oocytes indicates the SPC105R antibody detects a region between amino acids 124 and 473, while the region between 1 and 123 is not detected (Figure S 2).

### The role of SPC105R in homologous chromosome bi-orientation

To determine the effect of each *Spc105R* mutant on chromosome segregation, we examined homologous chromosome bi-orientation. Pairs of homologous chromosomes are bioriented in metaphase I when their centromeres are oriented towards opposite poles. To measure bi-orientation, we labelled three of the four *D. melanogaster* chromosomes, the X, 2^nd^, and 3^rd^, with fluorescent probes specific for each centromere. A bi-orientation defect was defined as both FISH signals oriented towards the same pole.

Chromosome bi-orientation in *Spc105R^B^, Spc105R^RNAi^*oocytes was similar to that of wild-type controls (Figure 2). *Spc105R^C^*, *Spc105R^RNAi^*oocytes had a high frequency of bi-orientation defects (61.9%), suggesting domains or sequences between amino acids 1 and 1284 regulate bi-orientation or error correction (Figure 2). A deletion of the large region containing ExxEED/E repeats, *Spc105R^ΔExxEED^, Spc105R^RNAi^* oocytes, did not have a significant frequency of bi-orientation defects. Deletion of this region was also fertile, but nondisjunction was observed among the progeny (Table 1). The difference is probably because the genetic assay is more sensitive, or there are errors in meiosis II. In contrast, *Spc105R^ΔM^ Spc105R^RNAi^*and *Spc105R^ΔMELT-KI^ Spc105R^RNAi^* oocytes had a high frequency of bi-orientation defects (45.5%) that was similar to *Spc105R^C^*. In addition, *Spc105R^ΔMELT-KI^* females, where the wild-type protein was present, were fertile and had significant levels of nondisjunction (Table 1). A lower but significant bi-orientation defect was observed in *Spc105R^ΔN^ Spc105R^RNAi^* oocytes (12.8%, p = 0.0023) (Figure 2). These results suggest that bi-orientation is primarily mediated by the region between amino acids 124 and 473, while the N-terminal domain has a minor contribution.

**Figure 2:**
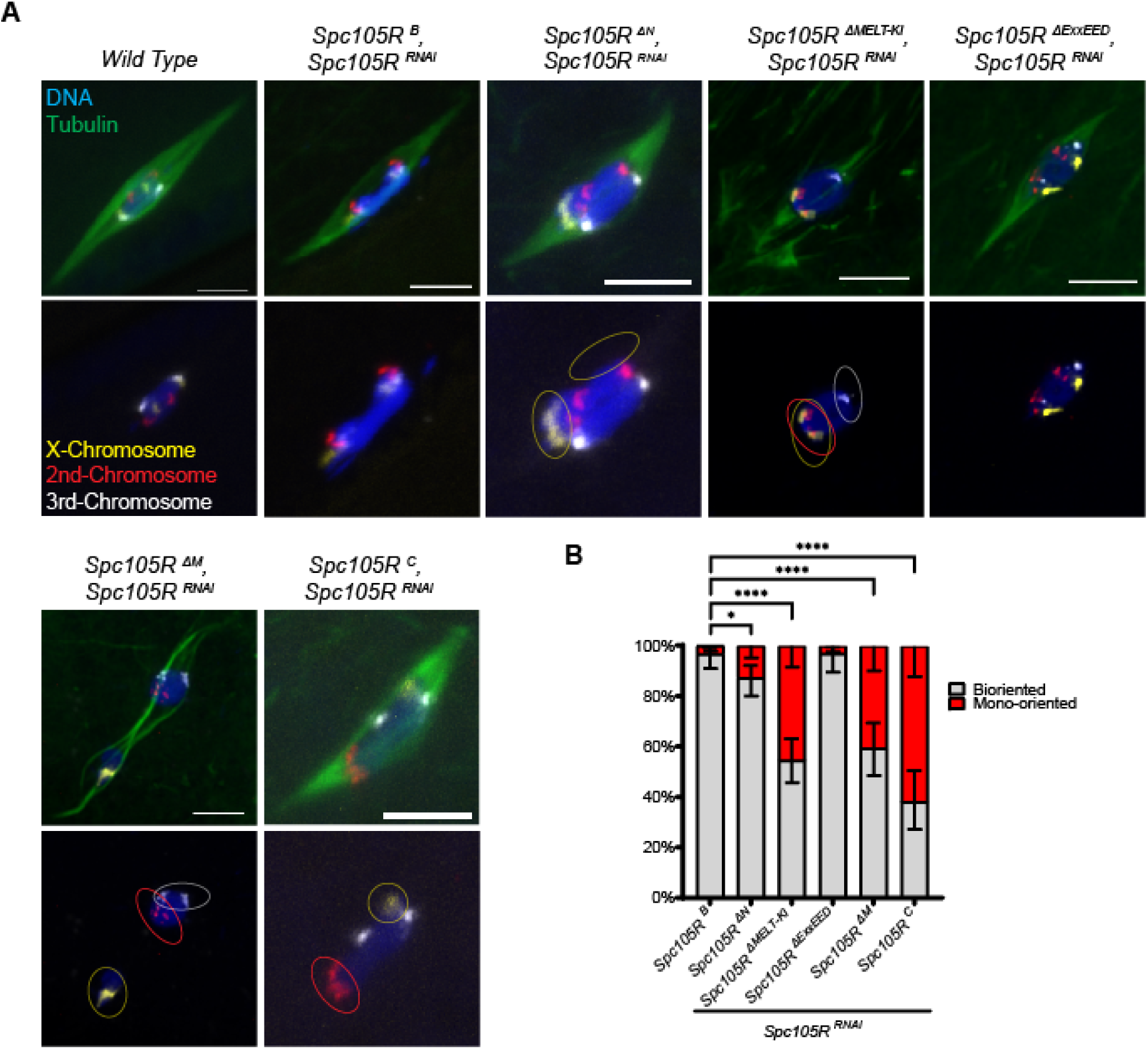
SPC105R domains required for homologous chromosome bi-orientation. A) Confocal images of *Spc105R^RNAi^*oocytes expressing the indicated transgene. Merged images show DNA (blue), Tubulin (green), the X-Chromosome (Yellow), the second chromosome (Red), and the third chromosome (white). DNA and FISH probes are shown in a separate channel. Scale bars are 5 μm. B) Quantification for percent of chromosome mono-orientation: *Spc105R ^B^*, *Spc105R^RNAi^* oocytes (n=93), *Spc105R ^ΔN^*, *Spc105R^RNAi^* oocytes (n=117), *Spc105R ^ΔExxEED^*, *Spc105R^RNAi^* oocytes (n=65), *Spc105R ^ΔM^*, *Spc105R^RNAi^*oocytes (n=81), *Spc105R ^ΔMELT-KI^*, *Spc105R^RNAi^* oocytes (n=123), *Spc105R ^C^*, *Spc105R^RNAi^*oocytes (n=63). Significance in frequency of mono-orientation om oocytes determined by Fisher’s exact test, with * = p-values < 0.01, **** = p-value < 0.0001.

### The C-terminal domain of SPC105R is sufficient for outer kinetochore assembly

The C-terminal domain of SPC105R, beginning after the last ExxEED/E repeat, was previously defined in mitotic cells as being capable of organizing the kinetochore (Schittenhelm et al., 2009). This KT-binding domain of SPC105R contains coiled-coil motifs which are suspected to drive its localization to the kinetochore and facilitate recruitment of other kinetochore proteins (Liu et al., 2016). In *Spc105R^C^*, *Spc105R^RNAi^*oocytes, we observed kinetochore localization of SPC105R^C^ using an antibody to the MYC epitope (Figure S 1C), showing this domain is sufficient for SPC105R recruitment in oocytes. However, when SPC105R^C^ was expressed in the presence of wild-type protein, both proteins were detected at the kinetochore (Figure S 2C). This may explain why expression of *Spc105R^C^*, and some other mutants, had a dominant phenotype and caused sterility even in the presence of wild-type SPC105R protein (Table 1).

In *Spc105R^RNAi^* oocytes, NDC80 does not localize to the kinetochore (Radford et al., 2015) (Figure 3A). In contrast, NDC80 localized to the kinetochores in *Spc105R^C^*, *Spc105R^RNAi^* oocytes, confirming that the C-terminal domain of SPC105R is sufficient to build a meiotic kinetochore (Figure 3A,B). Although *Spc105R^C^ Spc105R^RNAi^* oocytes could recruit NDC80 and assemble a kinetochore, the organization of the chromosomes and spindle was abnormal. In wild-type oocytes, all chromosomes coalesce together in a single round or oval karyosome. In *Spc105R^C^ Spc105R^RNAi^* oocytes, the karyosomes were often elongated and fragmented (Figure 3A, C), and contained a mix of end-on (Figure 4A and C, Figure S 1) and lateral (Figure 3A and C, Figure 4E) KT-MT attachments. The karyosome phenotype could be the result of creating a kinetochore that lacks key regulatory components recruited by the rest of SPC105R, resulting in unregulated microtubule attachments and causing the karyosome to elongate or fragment.

**Figure 3:**
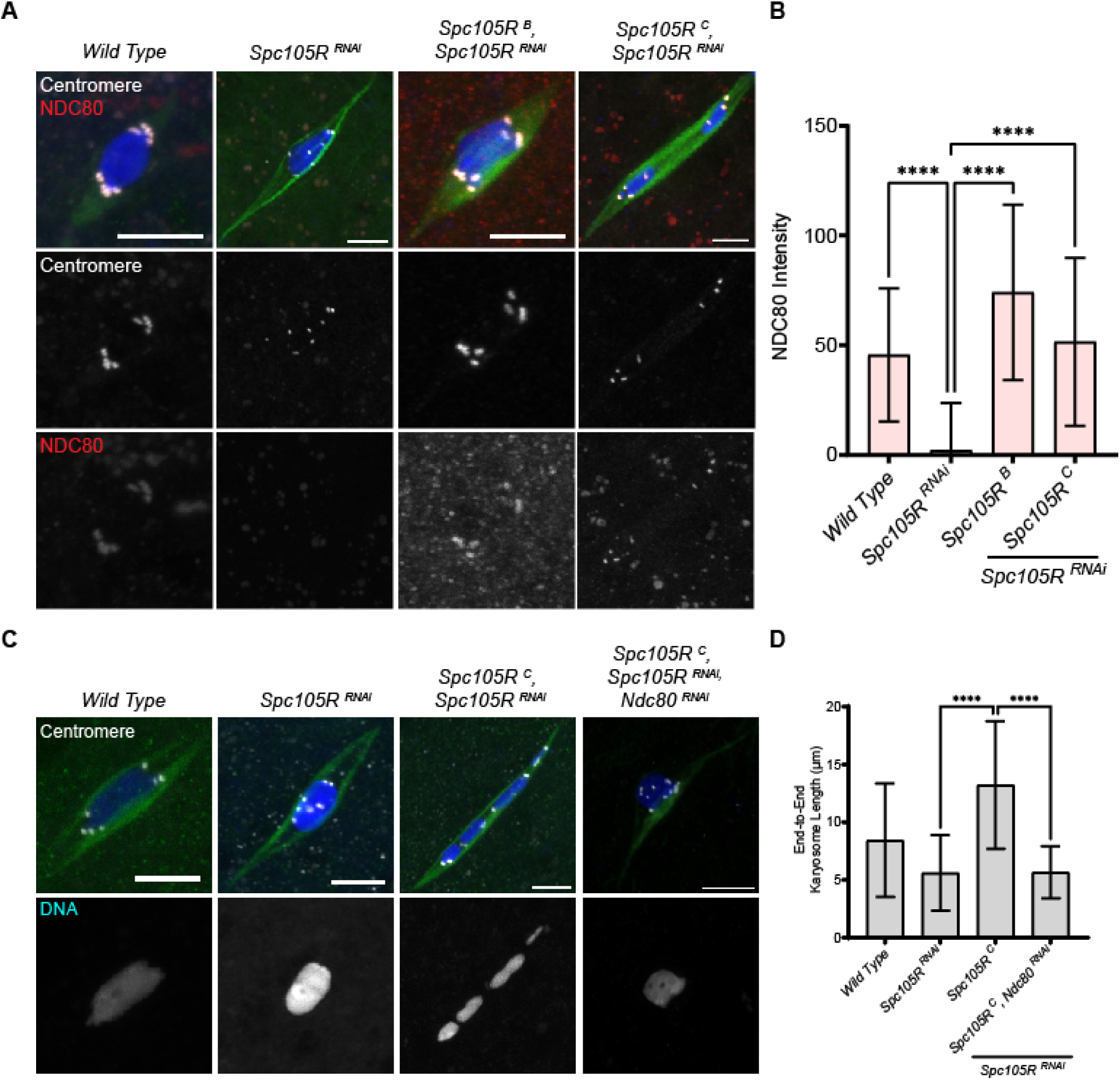
Spc105R’s C-Terminal Domain recruits NDC80. The N-terminal 1284 amino acids of SPC105R is deleted in *Spc105R ^C^*, retaining only the last 676 amino acids (Figure 1). A) Confocal images of *Spc105R^RNAi^*oocytes expressing the indicated transgene, with DNA (blue), Tubulin (green), centromeres (white), and the kinetochore protein NDC80 (red). Centromeres and NDC80 are shown in separate channels. Scale bars are 5 μm. B) Quantification of NDC80 intensity at the centromeres in w*ild-type* (n=68), *Spc105R*^RNAi^ (n=77), *Spc105R^B^*, *Spc105R*^RNAi^ (n=131), and *Spc105R^C^*, *Spc105R*^RNAi^ (n=107) oocytes. Error bars indicate standard deviation and **** = p-value < 0.0001 by a one-sided unpaired t-test. C) Confocal images of *Spc105R*^RNAi^ oocytes expressing the indicated transgene, with DNA (blue), Tubulin (green), and centromeres (white). DNA is shown in separate channels. Scale bars are 5 μm. D) Quantification of end-to-end chromosome length of *wild-type* (n=34), *Spc105R* ^RNAi^ (n=12), *Spc105R^C^*, *Spc105R*^RNAi^ (n=22), and *Spc105R ^C^*, *Spc105R*^RNAi^, *Ndc80*^RNAi^ (n=30) oocytes. Error bars indicate standard deviation, and *** = p-value < 0.001, **** = p-value < 0.0001, ns = not significant by a one-sided unpaired t-test.

**Figure 4:**
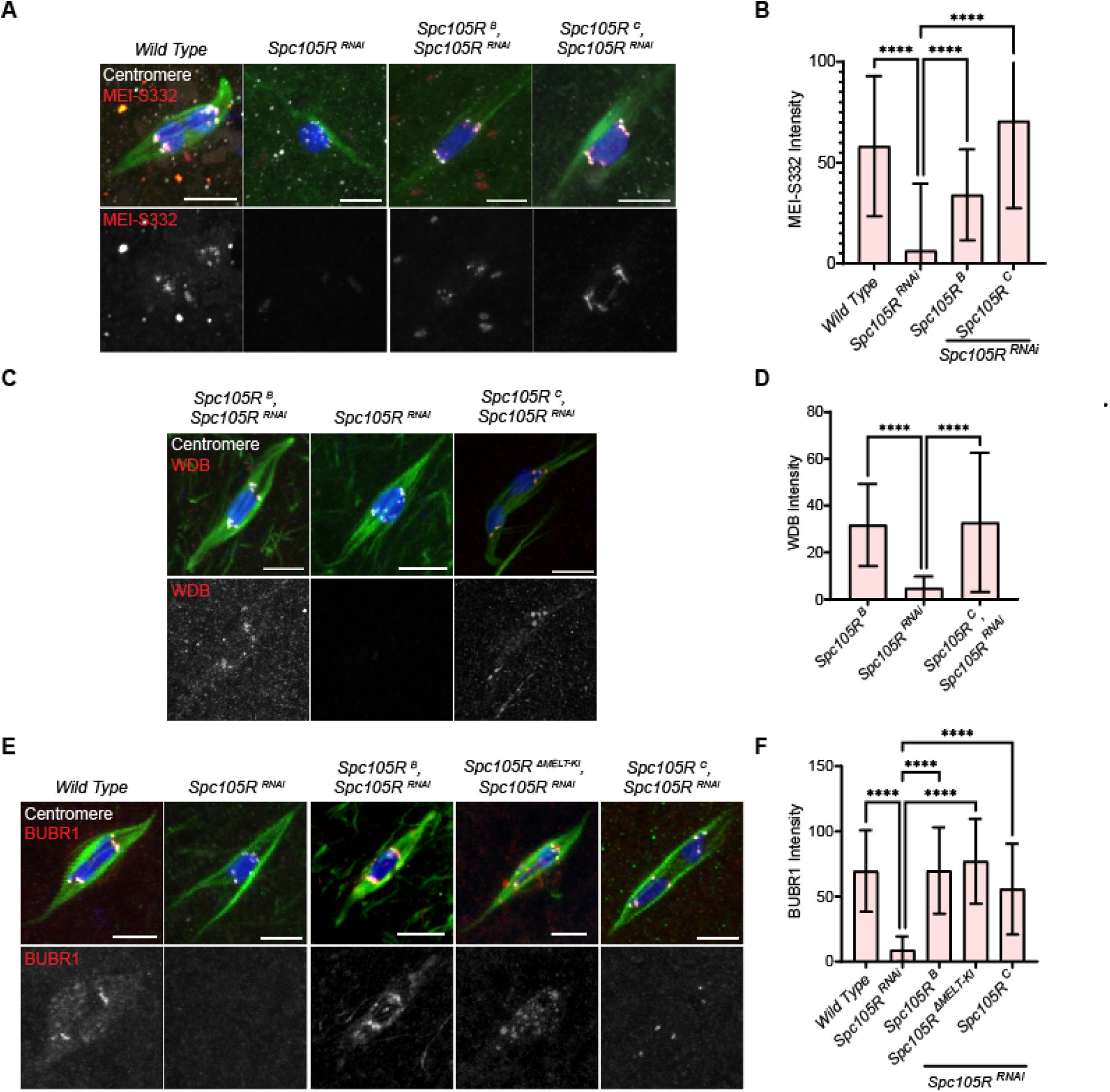
Spc105R’s C-terminal domain is required for centromeric cohesion protection in meiosis I. A) Confocal images of *Spc105R^RNAi^*oocytes expressing the indicated transgene with DNA (blue), Tubulin (green), centromeres (white), and MEI-S332 (red) and shown as a separate channel. Scale bars are 5 μm. B) Quantification of MEI-S332 intensity at the centromeres for w*ild-type* (n=71), *Spc105R*^RNAi^ (n=202), *Spc105R^B^*, *Spc105R*^RNAi^ (n=53) and *Spc105R^C^*, *Spc105R*^RNAi^ (n=114) oocytes. Error bars indicate standard deviation, and **** = p-value < 0.0001by one-sided unpaired t-test. C) Confocal images of *Spc105R*^RNAi^ oocytes expressing the indicated transgene, with DNA (blue), Tubulin (green), centromeres (white), and the PP2A subunit HA-WDB (red), also in a separate channel. D) Quantification of WDB intensity at the centromeres in *Spc105R^B^*, *Spc105R*^RNAi^ (n=134), *Spc105R*^RNAi^ (n=189) and *Spc105R^C^ Spc105R*^RNAi^ (n=155). Error bars indicate standard deviation and one-sided unpaired t-test show **** = p-value < 0.0001. E) Confocal images of *Spc105R*^RNAi^ oocytes expressing the indicated transgene, with DNA (blue), Tubulin (green), centromeres (white), and GFP-BUBR1 (red) and in a separate channel. F) Quantification of BUBR1 intensity at the centromeres, in *wild-type* (n=247), *Spc105R*^RNAi^ (n=153), *Spc105R^B^*, *Spc105R*^RNA^ (n=67) *Spc105R^C^*, *Spc105R*^RNAi^ (n =149) and *Spc105R^MELT-KI^*, *Spc105R*^RNAi^ (n =89) oocytes. Error bars indicate standard deviation and significance is shown by a one-sided unpaired t-test **** = p-value < 0.0001.

To determine if the karyosome elongation phenotype was caused by the unregulated formation of end-on KT-MT attachments, we examined *Spc105R^C^*, *Spc105R^RNAi^* oocytes also depleted for NDC80. The rationale for this experiment was that end-on attachments do not form in *Ndc80^RNAi^* oocytes (Radford et al., 2015). In *Spc105R^C^*, *Spc105R^RNAi^*, *Ndc80^RNAi^*oocytes, the karyosomes were spherical and shorter in length compared to *Spc105R^C^*, *Spc105R^RNAi^* oocytes (Figure 3C, D). Because NDC80 depletion suppressed the *Spc105R^C^ Spc105R^RNAi^* phenotype, it is likely that unregulated NDC80-dependent end-on attachments were responsible for the karyosome elongation and separation observed in *Spc105R^C^*, *Spc105R^RNAi^* oocytes.

### The C-terminal domain of SPC105R recruits proteins for maintaining cohesion

Co-orientation is the process to ensure sister kinetochores fuse and orient to the same pole in meiosis I (Watanabe, 2012). A co-orientation defect can result in merotelic attachments and errors in homologous chromosomes segregation. We previously showed that two mechanisms maintain co-orientation: sister centromere cohesion and the regulation of end-on attachment stability (Wang et al., 2019). In that study, we showed that SPC105R is required to prevent premature centromeric cohesion loss.

Centromeric localization of cohesion protection protein MEI-S332, the *Drosophila* orthologue of Shugoshin (SGO), depends on SPC105R (Figure 4A,B) (Wang et al., 2019). In *Spc105R^C^, Spc105R^RNAi^* oocytes, MEI-S332 was localized to the kinetochore. These results show that the C-terminal domain of SPC105R recruits MEI-S332 to the kinetochore. SGO recruits PP2A-B56, which maintains cohesion by dephosphorylating cohesin subunits and preventing Separase from cleaving the Kleisin subunit (Gutierrez-Caballero et al., 2012; Marston, 2015; Wassmann, 2013). We previously observed a reduction in the localization of the PP2A-B56 subunit WDB in a *mei-S332* mutant (Jang et al., 2021). Consistent with this result, WDB recruitment to the kinetochores was abolished by *Spc105R^RNAi^* but was present in *Spc105R^C^, Spc105R^RNAi^* oocytes (Figure 4C,D). These results show that the C-terminal domain of SPC105R is sufficient for the recruitment of PP2A-B56, and thus helps maintain sister-centromere cohesion during meiosis I.

Along with MEI-S332, BUBR1 is required for PP2A-B56 localization to the kinetochore (Jang et al., 2021). Given the C-Terminal domain was sufficient to recruit MEI-S332 and PP2A-B56, we tested if BUBR1 was recruited to the kinetochore by the C-Terminal domain of SPC105R. BUBR1 localization was absent in *Spc105R^RNAi^* oocytes. In contrast, BUBR1 localized to the kinetochores in *Spc105R^C^ Spc105R^RNAi^* oocytes (Figure 4E, F). Thus, the C-terminal domain of SPC105R is sufficient to recruit BubR1, MEI-S332, and PP2A-B56.

In mouse oocytes, MPS1 and BUB1 have been shown to recruit SGO2 (El Yakoubi et al., 2017). However, *Bub1^RNAi^* oocytes did not have a significant defect in MEI-S332 localization. (Figure S 3A,B), consistent with the observation that *Bub1^RNAi^* females were fertile. However, we found that MEI-S332 localization was reduced in *BubR1^RNAi^*and *Bub1^RNAi^, BubR1^RNAi^* oocytes and these females were sterile (Figure S 3A, B). Thus, BUBR1 is required to recruit MEI-S332, with possibly a minor contribution from BUB1.

Loss of cohesion can be observed by an increase in the number of centromere foci. In wild-type *Drosophila* meiosis I oocytes, 8 centromere foci representing the four *Drosophila* chromosomes are expected because the sister centromeres are fused. A defect in cohesion results in 9 to 16 centromere foci. The number of centromere foci in *Spc105R^C^ Spc105R^RNAi^* oocytes was similar to *Spc105R^B^ Spc105R^RNAi^* oocytes (Figure 5A,B), consistent with the conclusion that the C-terminal domain recruits cohesion protection proteins. *Bub1^RNAi^* and *BubR1^RNAi^*oocytes had significant defects in sister centromere cohesion, (Figure S 3C). In addition, consistent with redundancy, *Bub1^RNAi^*, *BubR1^RNAi^*oocytes had a more severe co-orientation defect than either single RNAi (10.7 centromere foci in *Bub1, BubR1 RNAi* oocytes compared to 8.5 in *Bub1* and 9.3 in *BubR1* RNAi oocytes). These results suggest that BUB1 and BUBR1 have partially overlapping functions in recruiting centromeric cohesion protection proteins.

**Figure 5:**
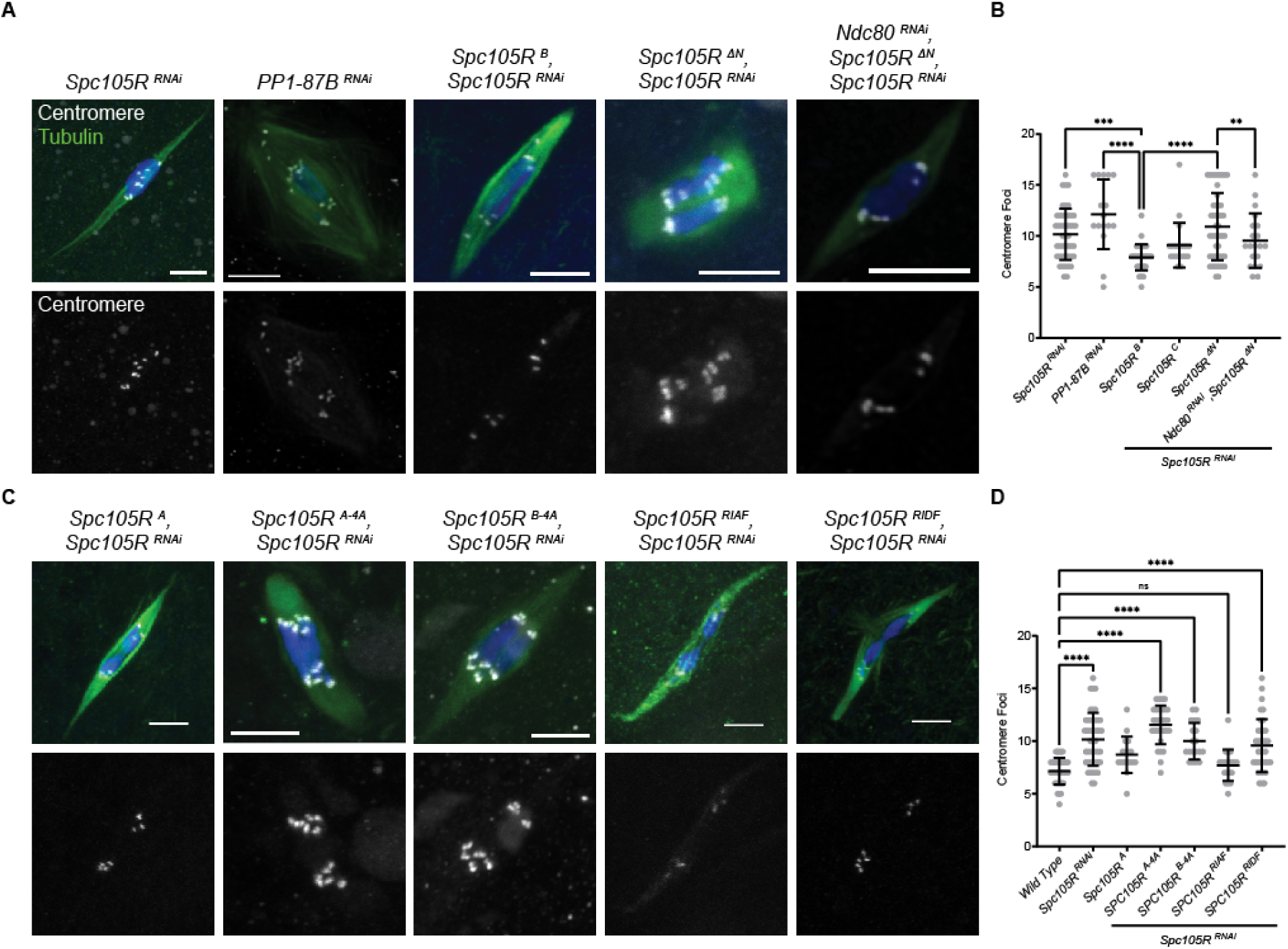
SPC105R N-terminal domain interacts with PP1 to maintain sister centromere co-orientation. Mutants with a deletion of the N-terminal domain (*Spc105R^ΔN^*) or mutations in the SLRK and RISF sequences are shown in Figure S 4A. A) Confocal images of *PP1-87B^RNAi^* or *Spc105R ^ΔN^* oocytes. Merged images show DNA (blue), Tubulin (green), and centromeres (white). Centromeres are shown in a separate channel. Scale bars are 5 μm. B) Quantification of centromere foci for images in A. One-sided unpaired t-test showed significance in number of foci. Sample sizes are: 46, 16, 37, 20, 48, 20, 16, in order of the graph. C) Confocal images of oocytes where the RISF site was changed to AAAA, RIAF or RIDF. D) Quantification of centromere foci for images in B. One-sided unpaired t-test showed significance in number of foci between all channels. Sample sizes are: 37, 46, 17, 29, 19, 23, 36, in order of the graph. Error bars indicate standard deviation, **** p < 0.0001, ** p = 0.001.

### The role of the N-terminal domain in mediating sister kinetochore co-orientation

Depletion of PP1-87B, one of three *Drosophila* PP1 alpha-type isoforms, causes a co-orientation defect that depends on the stabilization of end-on microtubule attachments (Wang et al., 2019) (Figure 5A, B). The SPC105R/KNL1 N-terminal domain has been shown to interact with PP1 (Bajaj et al., 2018). It is not known, however, if the co-orientation function of PP1 depends on its interaction with SPC105R. Therefore, we tested the hypothesis that the PP1 function in co-orientation is mediated by interactions with the N-terminal domain of SPC105R.

The co-orientation defect in *Spc105R^RNAi^* oocytes (11.6 centromere foci) was rescued by *Spc105R^B^* (Figure 5A, B) (8.2 centromere foci). A deletion of the entire N-terminal domain, *Spc105R^ΔN^ Spc105R^RNAi^* oocytes, had a significant increase in centromere foci, indicating a defect in co-orientation. The co-orientation defect in *PP1-87B^RNAi^*oocytes is dependent on end-on microtubule attachments (Wang et al., 2019). We used *Ndc80* RNAi to eliminate end-on attachments and found that the number of centromere foci was not elevated in *Ndc80^RNAi^*, *Spc105R^ΔN^*, *Spc105R^RNAi^* oocytes, and was similar to *Spc105R^B^ Spc105R^RNAi^*oocytes (Figure 5A, B). The suppression of the centromere foci phenotype is consistent with the conclusion that the co-orientation defect in *Spc105R^ΔN^*oocytes depends on end-on attachments.

The N-terminal domain of SPC105R contains two conserved Aurora B/PP1 interaction motifs: SSLRK and RISF. We mutated each site and counted the number of centromere foci to determine if there was a co-orientation defect (Figure S 4A). In *Spc105R^A^, Spc105R^RNAi^* oocytes, in which the serines in the SSLRK domain are absent, the number of centromere foci was not significantly changed compared to control wild-type oocytes (Figure 5C, D). In contrast, *Spc105R^A-4A^,* and *Spc105R^B-4A^*, *Spc105R^RNAi^*oocytes, in which RISF of either the A or B isoforms was changed to AAAA, had a significantly elevated number of centromere foci compared to wild-type oocytes, and were similar to the number of centromere foci in *Spc105R^ΔN^*and *PP1-87B*^RNAi^ oocytes. These results suggest that the RISF motif regulates centromere co-orientation. To test the effects of phosphorylation and PP1 binding on co-orientation, we made phosphomimetic (RIDF) and phosphodefective (RIAF) mutants. We observed that *Spc105R^RIDF^, Spc105R^RNAi^*, but not *Spc105R^RIAF^, Spc105R^RNAi^ oocytes*, had significantly increased centromere foci (Figure 5C, D). These results suggest that PP1 binding, which is predicted to be blocked by phosphorylation or the RIDF mutation, but not the RIAF mutation (Bajaj et al., 2018), may be the main cause of the co-orientation defect.

*PP1-87B^RNAi^* oocytes are also characterized by disorganization of the meiotic chromosomes (Wang et al., 2019). While wild-type oocytes chromosomes cluster together within a single karyosome, *PP1-87B^RNAi^* oocytes often have chromosomes separated into multiple groups. This phenotype was also observed in *Spc105R^ΔN^ Spc105R^RNAi^*and *Spc105R^RIDF^*, *Spc105R^RNAi^* oocytes, but not *Spc105R^RIAF^, Spc105R^RNAi^* oocytes (Figure S 4B, C). In addition, *Spc105R*^RIDF^, *Spc105R^RNAi^* oocytes had unique phenotypes, including tripolar and frayed spindles and evidence of enhanced interactions with the chromosome passenger complex (CPC). The CPC component INCENP is normally located on the central spindle of wild-type oocytes. In *Spc105R*^RIDF^, *Spc105R^RNAi^* oocytes, INCENP also localized at the kinetochores (Figure S 4B).

The *RIDF* mutant was not viable and sterile while the *RIAF* mutant was viable and fertile (Figure S 4D, Table 2). Furthermore, we did not observe significant defects in chromosome bi-orientation in the N-terminal domain mutants (Figure S 5A, B). Thus, phosphorylation (and PP1 binding ability) of SSLRK and RISF are implicated in regulating the metaphase I arrest, but not the bi-orientation of homologous chromosomes during meiosis I.

**Table 2:** Fertility and nondisjunction in *Spc105R* N-terminal and MELT-KI mutant females.

| *Spc105R<sup>RNAi</sup>* + | NDJ % | Total |
| --- | --- | --- |
| *Spc105R<sup>B</sup>* | 0.1 | 2347 |
| *Spc105R<sup>B-4A</sup>* | -- | sterile |
| *Spc105R<sup>A-4A</sup>* | -- | sterile |
| *Spc105R<sup>ΔKI</sup>* | 4.6 **** | 784 |
| *Spc105R<sup>ΔMELT</sup>* | -- | sterile |
| *SPC105R<sup>RIAF</sup>* | 0.0 | 322 |
| *Spc105R<sup>RIDF</sup>* | -- | sterile |
| No RNAi |  |  |
| *Spc105R<sup>B</sup>* | 0.5 | 2688 |
| *Spc105R<sup>B-4A</sup>* | -- | sterile |
| *Spc105R<sup>A-4A</sup>* | 0.4 | 463 |
| *Spc105R<sup>ΔKI</sup>* | 3.5 *** | 544 |
| *Spc105R<sup>ΔMELT</sup>* | 1.2 | 276 |
| *Spc105R<sup>RIAF</sup>* | 0.6 | 708 |
| *Spc105R<sup>RIDF</sup>* | -- | sterile |

Females expressing the indicated transgene were either in a *Spc105R<sup>RNAi</sup>* or wild-type background. Fishers exact test when compared to *Spc105R<sup>B</sup>*, \*\*\* = p = 0.001, \*\*\*\* = p < 0.0001. NDJ = nondisjunction

### The role of SPC105R regulatory domains in microtubule attachments

We previously showed that *Spc105R^RNAi^* oocytes have a more severe defect in microtubule attachments than *Ndc80* RNAi oocytes (Radford et al., 2015). These studies showed that NDC80 is required for end-on microtubule attachments while SPC105R is sufficient to mediate lateral microtubule attachments. However, how SPC105R mediates lateral attachments is not known. Therefore, we used two assays to identify the domain(s) of SPC105R that mediate interactions with microtubules.

The first assay for lateral attachments involved measuring the distance between each kinetochore and the closest microtubules. *Spc105R^RNAi^* oocytes have a significantly larger average kinetochore-microtubule distance than wild-type or *Ndc80^RNAi^*oocytes, indicating they lack the lateral attachments present in *Ndc80^RNAi^*oocytes (Figure S 6). All of the *Spc105R* mutants form end-on attachments through recruitment of NDC80 by the C-terminal domain. Therefore, to analyze microtubule interactions, each *Spc105R* mutant was analyzed in an *Ndc80* RNAi background. We predicted that in the absence of both NDC80 and a microtubule binding domain of SPC105R, KT-MT distances would be larger than the distance observed in *Ndc80^RNAi^* oocytes, and similar to the distance observed in *Spc105R^RNAi^* oocytes. We found that the KT-MT distances in *Ndc80^RNAi^*, *Spc105R^C^, Spc105R^RNAi^* oocytes were similar to the distances in *Spc105R^RNAi^* oocytes and significantly larger than the KT-MT distances in *Ndc80^RNAi^* oocytes (Figure S 5). This indicates that the C-terminal domain of SPC105R does not mediate lateral attachments.

We next tested whether the N-terminal domain has a role in lateral attachments by performing this analysis on two N-terminal domain deletion mutants. We chose *Spc105R^ΔN^* because it deletes a region previously shown to interact with microtubules (Audett et al., 2022). We also chose *Spc105R^ΔMELT-KI^* because it showed severe defects in homolog bi-orientation. We found that the KT-MT distances of both mutants were significantly lower than *Spc105R^RNAi^* oocytes, suggesting these two mutants can interact with microtubules. However, the KT-MT distances in *Spc105R^ΔN^, Spc105R^RNAi^* and *Spc105R^ΔMELT-KI^*, *Spc105R^RNAi^*were greater than in *Ndc80^RNAi^* oocytes, suggesting a defect in microtubule attachments. Because these mutants were not as severe as *Spc105R^C^, Spc105R^RNAi^* oocytes, these results could be explained if multiple SPC105R domains mediate lateral attachments.

The second assay was based on an observation that lateral attachments are sufficient for chromosome movement. Oocytes lacking chiasma precociously enter anaphase (McKim et al., 1993). We used *mei-P22* mutants, which lack DSBs, to generate oocytes without chiasmata (Liu et al., 2002). In *mei-P22^-^, Ndc80^RNAi^* oocytes, precocious chromosome movement towards the spindle poles, as shown by an elongated spindle with a separated karyosome, was observed (Figure 6A). In contrast, chromosome movement towards opposite poles was significantly reduced in *mei-P22^-^, Spc105R^RNAi^* oocytes (Figure 6B). These results show that the SPC105R-dependent lateral attachments are sufficient for chromosome movement. We then used this assay to identify the SPC105R domain responsible for lateral attachments. We generated females expressing *Spc105R* mutants in a *mei-P22^-^, Spc105R^RNAi^*, *Ndc80^RNAi^*background. As expected, *Spc105R^C^* was defective in movement towards the poles (Figure 6A), suggesting a region between amino acids 1 and 1284 mediates chromosome movement. *Spc105R^ΔN^* was capable of movement towards the poles, showing that this domain was not required for microtubule interactions. *Spc105R^ΔMELT-KI^*, however, was defective in chromosome movement. These results suggest that the region including the MELT- and KI-like repeats between amino acids 123 and 473 are required for chromosome movement, consistent with a role in facilitating lateral attachments to microtubules.

**Figure 6:**
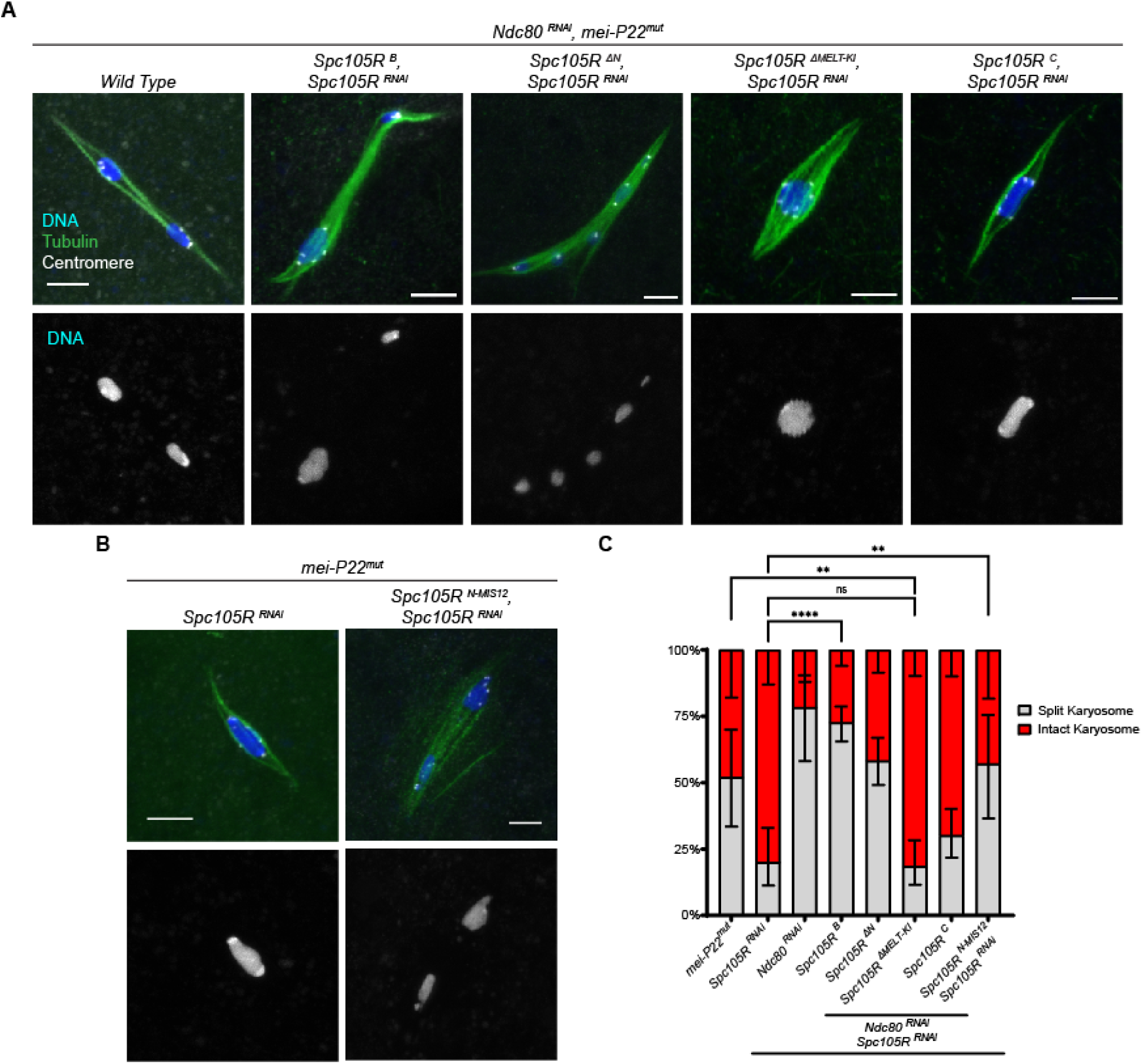
SPC105R regulates interactions with microtubules. A) Confocal images of oocytes in a *mei-P22* mutant (*mei-P22^mut^*) background. The first image is only *Ndc80 RNAi,* while the rest express *Ndc80^RNAi^*, *Spc105R^RNAi^*, and a *Spc105R* transgene. Due to the absence of crossing over, *mei-P22^mut^* mutant oocytes fail to arrest in metaphase, and precociously enter anaphase. B) Oocytes in *mei-P22^mut^, Spc105R* RNAi background. The second image is also expressing a *Spc105R-mis12* transgene. Merged images show DNA (blue), Tubulin (green), and centromeres (white). Chromosomes are shown in a separate channel. Scale bars are 5 μm. C) Quantification of the frequency of chromosome separation in oocytes. Sample sizes are: 25, 50, 23, 175, 115, 81, 83, 21, in order of the graph. Significance in frequency of oocytes entering precocious anaphase determined by Fisher’s exact test, with ** = p-value < 0.01, **** = p-value < 0.0001, and ns = not significant.

### Lateral attachments are not sufficient for bi-orientation

To test if SPC105R-mediated lateral attachments are sufficient for bi-orientation, we compared FISH results in *Ndc80^RNAi^*and *Spc105R^RNAi^* oocytes. The high frequency of bi-orientation errors in *Spc105R^RNAi^* oocytes was not surprising given the lack of KT-MT attachments (Figure S 5C,D). However, *Ndc80^RNAi^*oocytes were not significantly better, despite having lateral attachments. Thus, lateral attachments mediated by SPC105R are not sufficient for accurate bi-orientation of homologous chromosomes at meiosis I.

### Dissecting the MELT-KI region

To investigate the functions of the MELT-KI region, we created two mutants with a deletion of each domain. For the KI domain, we deleted amino acids 247-473 of SPC105R. In these *Spc105R^ΔKI^*, *Spc105R^RNAi^* oocytes, we did not observe a statistically significant difference in bi-orientation defects (15% mono-orientation) when compared with *Spc105R^B^*, *Spc105R^RNAi^* (Figure S 5). In addition, *Spc105R^ΔKI^*, *Spc105R^RNAi^* females were viable, fertile, and had a low, but significant, frequency of nondisjunction (Figure S 4D, Table 2). These data suggest that the KI region may not be required for most aspects of meiosis, particularly the regulation of bi-orientation (Figure S 5). To examine the function of the MELT-like repeats, we created a *Spc105R^ΔMELT^*mutant with a deletion of amino acids 123-246. *Spc105R^ΔMELT^*, *Spc105R^RNAi^* females were sterile, consistent with a defect in embryonic mitosis, but like the *ΔKI* mutant, were viable (Figure S 4, Table 2). Similarly, *Spc105R^ΔMELT^*, *Spc105R^RNAi^* oocytes had an insignificant increase in the frequency of bi-orientation defects (Figure S 5B). Thus, relative to the larger deletion of both domains (*Spc105R^ΔMELT-KI^*), the *Spc105R^ΔKI^* and *Spc105R^ΔMELT^* mutants had weaker phenotypes. These results suggest that the MELT and KI domains may function together or additively to facilitate homologous chromosome bi-orientation in meiosis and possibly also in mitosis.

### Testing the function of the N-terminal domain with a fusion to MIS12

To independently test if the domains of SPC105R not involved in kinetochore assembly are sufficient for microtubule interactions, we fused the N-terminal 1284 amino acids of SPC105R to MIS12. The fusion protein localized to centromeres in *Spc105RN-Mis12, Spc105R^RNAi^* oocytes (Figure 7A,B) but failed to recruit NDC80 (Figure 7C, D). In addition, we observed the staining of puncta on the spindle, suggesting that the SPC105RN-MIS12 complex detaches from the chromosomes, possibly in a multiprotein complex (Figure 7A). This effect was more severe in the presence of wild-type SPC105R, suggesting the interaction between SPC105RN-MIS12 and the centromeres is unstable when there is competition from the wild-type protein or in the presence of end-on attachments.

**Figure 7:**
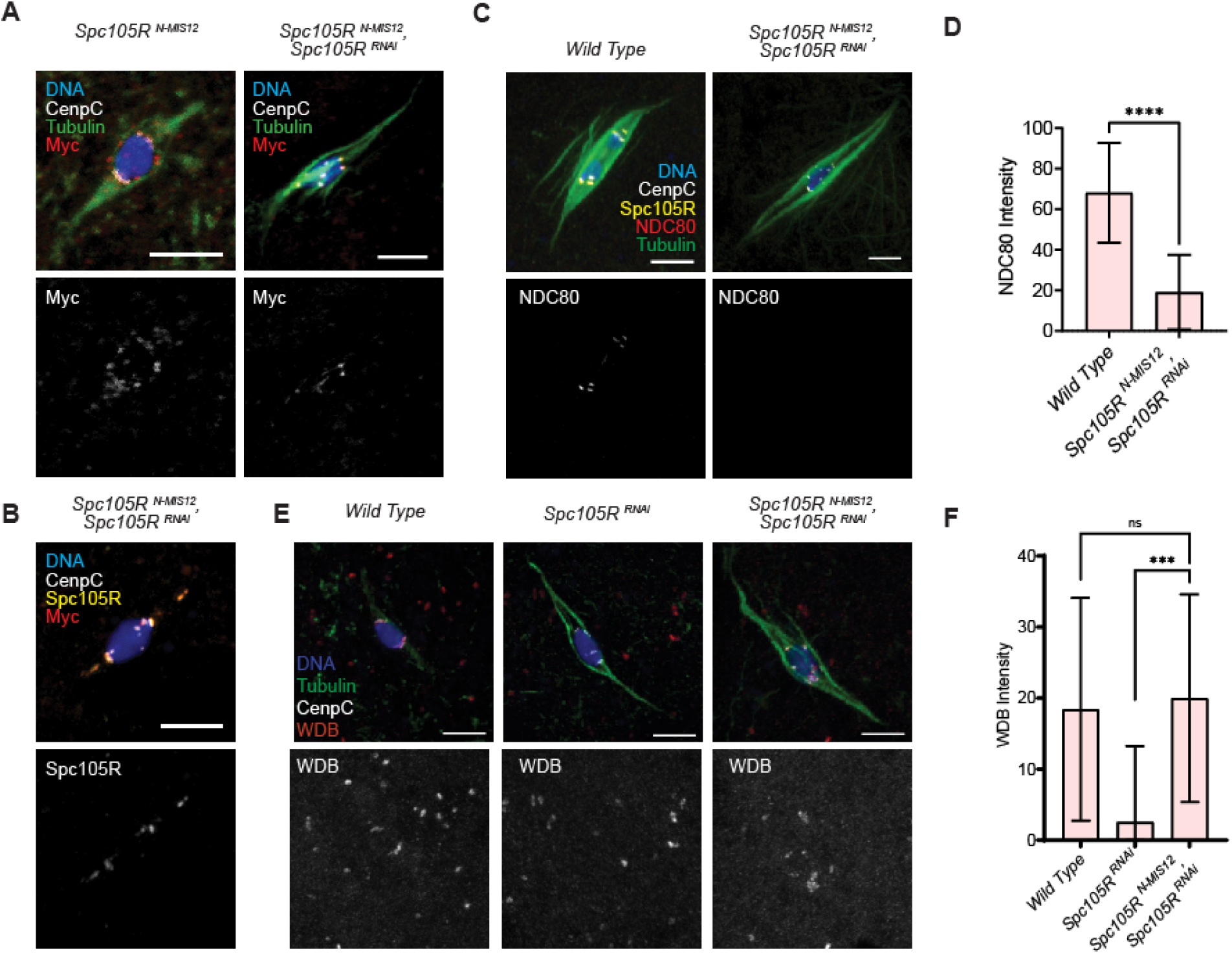
The N terminal domain of SPC105R fused to MIS12 A) Confocal images of stage 14 oocytes expressing *Spc105R^N-Mis12^,* either in the presence or absence of *Spc105R^RNAi^*. The fusion protein was detected using an antibody to the Myc tag (red), along with CENP-C (white), and tubulin (green). B) *Spc105R^N-MIS12^, Spc105R^RNAi^*oocyte, detected using both an antibody to the Myc tag (red) and SPC105R (yellow), along with CENP-C (white). C) Ndc80 (red) does not localize in *Spc105R^N-MIS12^, Spc105R^RNAi^* oocytes. D) Measurement of NDC80 intensity in wild-type (n=137) and *Spc105R ^N-MIS12^, Spc105R^RNAi^* (n=160) oocytes. Error bars indicate standard deviation, and one-sided unpaired t-test showed significance **** = p < 0.0001. E) WDB (red) localization in *Spc105R ^N-MIS12^, Spc105R^RNAi^* oocytes. F) Measurement of WDB intensity in wild-type (n=154), *Spc105R^RNAi^* (n=195), and *Spc105R^N-MIS12^, Spc105R^RNAi^* (n=183) oocytes. Error bars indicate standard deviation and one-sided unpaired t-test showed significance, *** = p < 0.001.

When expressed with the *mei-P22* mutant allele, *Spc105R-Mis12, Spc105R^RNAi^* oocytes showed chromosome movement towards the poles, suggesting the kinetochores make lateral attachments to the microtubules (Figure 6B, C). Given the failure to recruit NDC80, the microtubule interactions in these oocytes occur via SPC105R. These results confirm that the N-terminal domain of SPC105R, most likely the MELT-KI region, is sufficient for lateral microtubule interactions and chromosome movement.

We have shown that the C-terminal domain of SPC105R recruits BUBR1 and PP2A. However, the region of vertebrate KNL1 corresponding to MELT-KI in SPC105R has also been shown to recruit BUBR1 and PP2A (Caldas and DeLuca, 2014; Saurin, 2018). These two results raise the possibility that SPC105R has two BUBR1/ PP2A recruitment sites. To test this possibility, we stained *Spc105R-Mis12, Spc105R^RNAi^* oocytes for PP2A. These results showed that the *Spc105R-Mis12, Spc105R^RNAi^* oocytes were able to recruit PP2A (Figure 7E,F), and combined with experiments using the C-terminal domain (Figure 4), indicates that SPC105R does have two distinct sites for recruiting PP2A.

## Discussion

### A system for studying kinetochore function in Drosophila oocytes

SPC105R is at the center of a complex network of signals that regulate microtubule attachments and cell cycle progression (Saurin, 2018). The studies presented here were undertaken to investigate two functions critical for meiotic kinetochores of *Drosophila* oocytes. First, two mechanisms promote co-orientation of sister centromeres at meiosis I, cohesion protection and regulating the stability of microtubule attachments (Wang et al., 2019). Second, that SPC105R is sufficient for lateral microtubule attachments (Radford et al., 2015).

### Recruitment of the outer kinetochore and cohesion protection

The C-terminal domain of SPC105R is required for kinetochore assembly while the rest of the protein regulates microtubule attachments and the spindle assembly checkpoint. This is consistent with previous studies showing that the NDC80 complex (NDC80c) is recruited by the C-terminal domain of SPC105R/KNL1 in *Drosophila* mitotic cells (Schittenhelm et al., 2009). However, NDC80c can also be recruited by KNL1-independent mechanisms in vertebrate cells, such as the CENP-T pathway (Sridhar and Fukagawa, 2022). These mechanisms depend on proteins that have not been identified in *Drosophila*. Our results suggest that these other pathways have been lost and SPC105R is the only pathway to assemble NDC80c.

For mitotic cells, protection of cohesins in pericentric regions in critical. The sister kinetochores need to remain separate in order to biorient. Our results provide new insights into the mechanism of maintaining cohesins at meiosis I centromeres, which is important to fuse the sister kinetochores for co-orientation (Nasmyth, 2015; Watanabe, 2012). We previously found that SPC105R is required to maintain sister centromere cohesion (Jang et al., 2021; Wang et al., 2019). Here we have shown that the C-terminal domain of SPC105R recruits cohesion protection proteins BubR1, MEI-S332 and PP2A-B56 and is sufficient to maintain sister-centromere cohesion.

Cohesion is established during premeiotic S-phase while SPC105R, like other outer kinetochore proteins, is not recruited to the centromeres until prometaphase. Thus, during a prolonged prophase arrest, the *Drosophila* oocyte must protect centromere cohesion by a kinetochore-independent mechanism. CENP-C may have this role in *Drosophila* (Fellmeth et al., 2023), similar to the role of mammalian and yeast CENP-C in recruiting Moa1 or Meikin (Nasmyth, 2015; Watanabe, 2012). Moa1 and Meikin are not conserved, but *Drosophila* Matrimony may have this role by regulating Polo kinase, which is also required for cohesion maintenance in *Drosophila* (Bonner et al., 2013; Bonner et al., 2020; Kim et al., 2015).

We propose that in meiosis I, SPC105R recruitment of PP2A maintains meiosis I co-orientation by providing the cohesion protection that keeps the sister kinetochores fused as a single unit. PP2A may antagonize Polo kinase, which is required for cohesion loss in *Spc105R*-depleted oocytes (Wang et al., 2019) and could be responsible for the phosphorylation event that is required for Separase cleavage of cohesin (Keating et al., 2020). In some mitotic cell types, SGO is recruited to pericentric regions and depends on BUB1 phosphorylation of H2A (Kawashima et al., 2010; Miyazaki et al., 2017). While BUB1 is known to recruit SGO in mouse meiosis (El Yakoubi et al., 2017), *Drosophila* BUB1 knockdown has a surprisingly weak effect on meiosis and viability. BUBR1, on the other hand, had the strongest effect on co-orientation, and has been shown to be required for sister chromatid cohesion in *Drosophila* male and female meiosis (Malmanche et al., 2007). BUB1 may have a minor role in cohesion maintenance, based on the observation that the double knockdown of *Bub1* and *BubR1* had the most severe co-orientation defect.

Although MEI-332 localizes to metaphase I chromosomes, it is only required for sister chromatid cohesion in meiosis II (Kerrebrock et al., 1995; Tang et al., 1998; Wang et al., 2019). MEI-S332 may be redundant in meiosis I with Dalmatian (DMT), a Soronin orthologue, in recruiting PP2A-B56 (Jang et al., 2021; Yamada et al., 2017). We do not know whether SPC105R is required to recruit DMT, but the strong cohesion defect in *Spc105R* RNAi oocytes suggests this may be the case.

### The N-terminal domain, checkpoint control, and co-orientation

Depletion of PP1-87B in meiosis results in the separation of sister centromeres (a co-orientation defect) and the chromosomes at metaphase I (Wang et al., 2019). Furthermore, the N-terminal domain of SPC105R has been shown to interact with PP1 in a variety of species (Audett et al., 2022; Bajaj et al., 2018; Roy et al., 2019). We found that deletion of the N-terminal domain or mutations of the RISF motif resulted in a phenotype similar to loss of PP1. The increase in centromere foci observed in oocytes lacking PP1 depends on microtubule attachments and is independent of Separase (Wang et al., 2019). These results suggest that the SPC105R-PP1 interaction regulates the stability of microtubule attachments, and this is important for maintaining co-orientation.

As done by others in studying the checkpoint response (Rosenberg et al., 2011; Roy et al., 2019), we have characterized phospho-mimetic and -defective mutants to test the effects of Aurora B phosphorylation and PP1 localization on co-orientation. Co-orientation defects were found in mutants that were predicted to prevent PP1 binding to SPC105R (RIDF and AAAA). In contrast, a mutant preventing Aurora B phosphorylation (RIAF) had mild effects on meiosis, fertility and viability. These results suggest that a defect in PP1 binding, which is blocked by phosphorylation of RISF (Bajaj et al., 2018) or the RIDF mutation, may be the main cause of the co-orientation defect. This is consistent with the loss-of-function phenotype of PP1. In addition, the RIDF mutant phenotype showed increased CPC at the centromeres, suggesting PP1 and the CPC compete for binding to the RISF site.

The mechanism of the co-orientation and karyosome separation phenotypes is not known. PP1 binding to the RISF motif of SPC105R has been associated with satisfaction of the spindle assembly checkpoint but not error correction (Bajaj et al., 2018; Espeut et al., 2012; Rosenberg et al., 2011; Roy et al., 2019). Our results, specifically that *PP1-87B^RNAi^* (Wang et al., 2019) and the RISF mutant oocytes did not have biorientation defects, are consistent with these conclusions. In *Drosophila* mitotic cells, binding of PP1 to the N-terminal domain promotes SAC satisfaction (Audett et al., 2022). Our results suggest that the failure to recruit PP1 results in premature sister centromere and karyosome separation, which is opposite to the expected result if there was a failure to satisfy the SAC. Instead, metaphase I arrest may depend on some of the same signals that trigger anaphase in mitotic cells. This is consistent with the observation that lack of tension on the kinetochores results in premature anaphase (Jang et al., 1995; McKim et al., 1993). Reversal of SAC signaling on metaphase progression may be important for oocytes with natural arrest points. The signals for progression into anaphase in meiosis I may be different than in mitosis and PP1 binding to SPC105R appears to maintain the metaphase I arrest.

### Lateral attachments and bi-orientation

Oocytes depleted of *Ndc80* or *Spc105R* have a “hollow spindle” phenotype, being made entirely of interpolar microtubules. In addition, SPC105R is sufficient for lateral microtubule attachments (Radford et al., 2015). Lateral attachments rely on interpolar microtubules, which can promote bi-orientation in mitosis (Magidson et al., 2011; Renda et al., 2022) and meiotic cells (Jang et al., 2005; Kitajima et al., 2011). We have shown here that lateral attachments are also capable of poleward movement during meiosis, as is also the case in *Drosophila* mitotic cells (Feijão et al., 2013). Because lateral attachments precede end-on attachments, they could have a significant role in bi-orientation, for example by preceding and preventing precocious end-on attachments (Itoh et al., 2018b; Muscat et al., 2015). However, we observed that bi-orientation was defective in *Ndc80* RNAi oocytes which have lateral attachments. One explanation for this result is that lateral interactions in wild-type cells may involve the interpolar microtubules interacting with short NDC80-dependent kinetochores microtubules (Doodhi and Tanaka, 2022). The short kinetochore fibers would give the lateral interactions directionality and perhaps provide a mechanism for bi-orientation prior to making end-on attachments (Renda et al., 2022).

The MELT-KI region, or amino acids 124-473, has two interesting properties. First, it has a role in making lateral attachments. Second, the MELT-KI domain is more important than any other region for bi-orientation of homologous chromosomes. The *Drosophila* MELT-KI domain contains two components, the first half with the MELT-like motifs, and the second half with KI-like motifs. The presence of 1-3 MELT-like repeats upstream of the KI motifs is a conserved feature of KNL1/SPC105R-like proteins (Tromer et al., 2015). Indeed, a vertebrate MELT-KI module was defined as sufficient for SAC activity (Vleugel et al., 2013)(Zhang et al., 2014). In our experiments, deletion of either the MELT motifs or the KI like motifs was less severe than deleting both. Thus, these two components may be part of a single domain required for bi-orientation of homologous chromosomes, as well as other function in mitosis that contribute to fertility and viability.

With sequences that resemble MELT motifs, the MELT-KI region could interact with MPS1 and Aurora B kinase, proteins known to be required for accurate meiotic chromosome segregation (Colombié et al., 2008; Gilliland et al., 2007; Radford et al., 2012). We have also shown that the N-terminal half of SPC105R, possibly the MELT-KI region, recruits PP2A. This activity could regulate bi-orientation given that PP2A is required for conversion of lateral to end-on attachments in *Drosophila* oocytes (Jang et al., 2021) and in mitotic cells (Caldas and DeLuca, 2014; Keating et al., 2020). It is also a region that may recruit the motor protein CENP-E, which has also been shown to be required for meiotic chromosome bi-orientation in *Drosophila* oocytes (Radford et al., 2015) and conversion from lateral to end-on attachments (Huang et al., 2019; Shrestha and Draviam, 2013). Finally, it has been shown that this region recruits the RZZ complex (McGory et al., 2024), which has been proposed to inhibit end-on attachments (Barbosa et al., 2022; Cheerambathur et al., 2013). To understand the mechanism of accurate segregation of chromosomes at meiosis I, it will be important to identify the proteins that interact with this region that are required for bi-orientation.

## Materials and Methods

### Drosophila genetics and qRT-PCR

*Drosophila* stocks and crosses were kept at 25 degrees Celsius and maintained on standard media. Many stocks used for experiments were acquired from the Bloomington Stock Center or from the Harvard Transgenic RNAi Project (TRiP). A list of stocks and their origins is in Table 3. In most experiments, expression of *UASP* transgenes was induced using *P{w[+mC]=matalpha4-GAL-VP16}V37* (referred to as *matα*). Expression begins during late pachytene in prophase (stage 1), which is after premeiotic DNA replication, and continues through late prophase until the oocyte matures (stage 14) (Radford et al., 2012; Sugimura and Lilly, 2006).

**Table 3.**
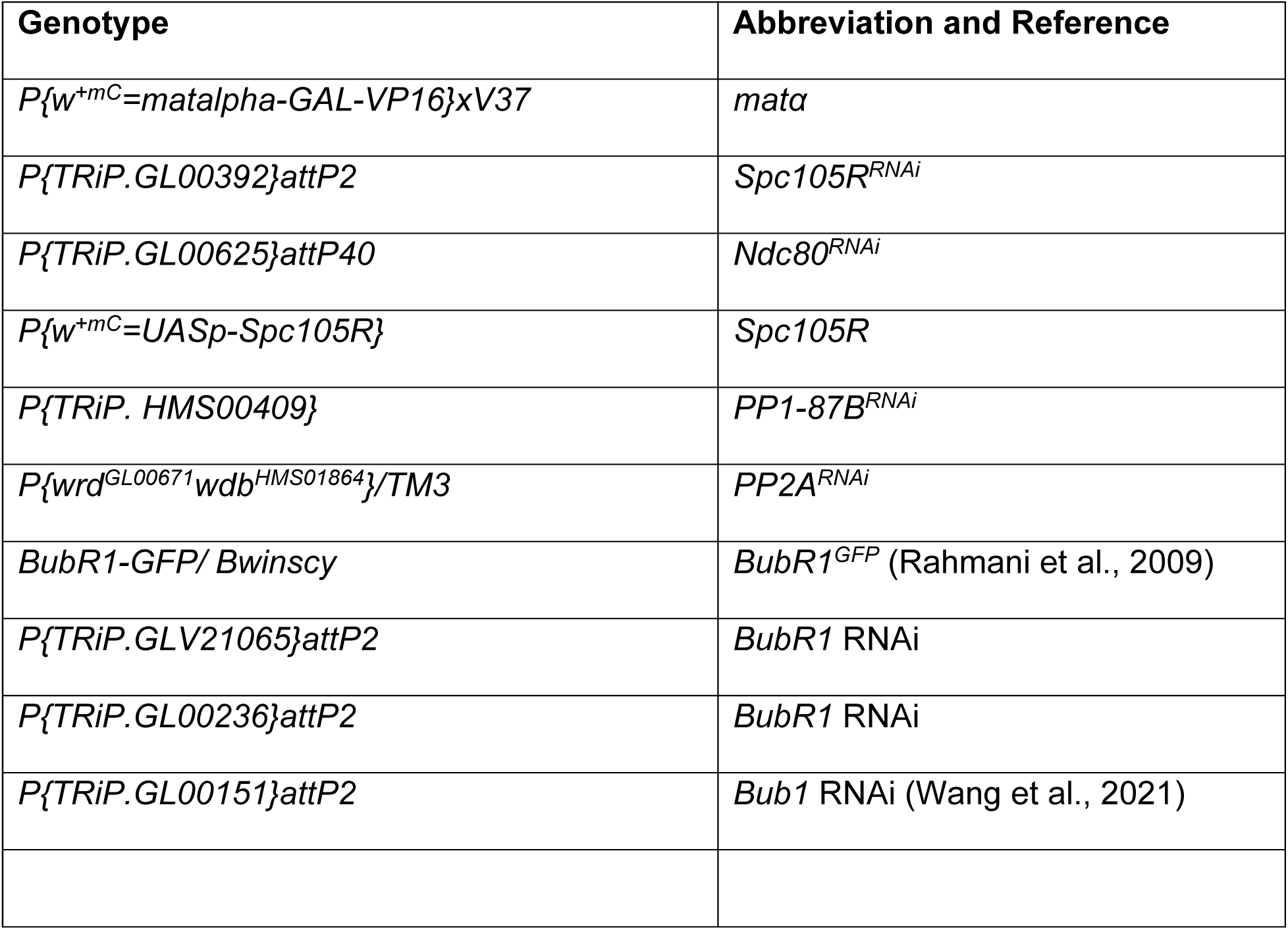
*Drosophila* transgenes used in this work.

To determine the effectiveness of shRNAs in depleting mRNA, total RNA was extracted from oocytes using TRIzol® Reagent (Life Technologies) and reverse transcribed into cDNA using the High Capacity cDNA Reverse Transcription Kit (Applied Biosystems). TaqMan® Gene Expression Assays (Life Technologies) were used for qPCR on a StepOnePlus™ (Life Technologies) real-time PCR system. The shRNA line and percent of wild-type mRNA levels remaining were: *Ndc80 GL00625* = 6%*, Spc105R GL00392* = 13% (Radford et al., 2015)*, Bub1 GL00151* = 2% (Wang et al., 2021), *BubR1 GLV21065* = 22%, and *BubR1 GL00236*= 27%.

### Generation of SPC105R Transgenes

The *Spc105R* transgenes were generated from the fully sequenced cDNA clone IP22012, which corresponds to the A splice form (*Spc105R^A^*). The 5’UTR and coding region with the stop codon removed were cloned into the pENTR4 vector. The transgene was made resistant to RNAi by making silent mutations in the sequences complementary to *GL00392.* The sequence in *Spc105R* targeted by GL00392, AAGGAGGATAGTTTAGCTAGA, was changed to AAGGAGGACAGCCTGGCCCGC.

The difference between the A and B forms is the failure to splice the first intron. The B splice form (*Spc105R^B^*) was generated using site directed mutagenesis to delete the first intron from the A form (NEB BaseChanger Kit). All mutants were generated using site specific mutagenesis of the A- or B-form *Spc105R* coding region in the pENTR4 vector. Wild-type and mutant clones were then subcloned into *pPWM* using the Clonase reaction (Invitrogen), resulting in fusion of the coding region to the 6 copies of the myc tag in the 3’ end. Constructs of *Spc105R* with deletions or mutations of these domains were injected into *Drosophila melanogaster* embryos (Model System Genomics or BestGene). The linkage of the transgenes was determined and insertions on the 3^rd^ chromosome were recombined onto the same chromosome as the shRNA *GL00392*.

### Generation of Spc105R mutants, fertility and viability

The genetic and cytological phenotypes of each mutant were studied by simultaneous tissue-specific knockdown of endogenous *Spc105R* by RNAi (*Spc105R^RNAi^* oocytes) and expression of a *Spc105R* transgene. For example, for control experiments we generated females carrying the two UAS transgenes, the *shRNA* and the wild-type *Spc105R*, and *matα*, which will be referred to as *Spc105R^B^, Spc105R^RNAi^*oocytes. Dominant phenotypes were examined by expressing mutant transgene in the absence of the *shRNA*.

In order to test for somatic functions such as mitosis, we used *P{w[+mC]=tubP-GAL4}LL7* (referred to as *Tub:GAL4*), which promotes expression in all tissues. Viability of the mutant *Drosophila* was measured by crossing *Spc105R^RNAi^*males to *Tub:GAL4/TM3, Sb* females. Progeny that inherited the *Tub:GAL4* and *Spc105R* transgenes were expected to survive if the mutant was functional in mitosis. The Sb progeny (*Spc105R, RNAi /TM3, Sb*) served as the control because they did not express the transgenes and were expected to survive.

To measure fertility and nondisjunction of the X chromosome, *Spc105R^B^, Spc105R^RNAi^* virgin females were crossed to males with a dominant *Bar* mutation on the Y chromosome (*yw/B^S^Y*). Unique phenotypes for the normal segregation of chromosomes (XX) or (XY), and nondisjunction of the X chromosome, (XO, XXY, XXX, and YO), were scored. XO and XXY progeny are viable whereas XXX and YO are lethal. To calculate nondisjunction rates while taking into account the lethal genotypes, the following equation was used: ^#^ *^flies^ ^with^ ^nondisjunction^ ^x 2^*. (# *flies with nondisjunction x 2) + (*# *of normal flies)*

### Immunofluorescence and microscopy

Oocytes in the 14th stage of oocyte development were collected from 2-to 3-day-old, yeast-fed non-virgin females (Radford and McKim, 2016). These flies were then ground in 1x modified Robb’s buffer and filtered through meshes to isolate the stage 14 oocytes. The oocytes were fixed with 5% formaldehyde and heptane before being rinsed in 1X PBS. The oocytes were then rolled between glass slides to remove the membranes and incubated for two hours in PBS/1% Triton X-100 to make them permeable to antibody staining. Afterwards, oocytes were washed in PBS/0.05% Triton X-100 and blocked in PTB (0.5% BSA and 0.1% Tween-20 in 1X PBS) for 1 hour.

Subsequently, primary antibodies were added to the oocytes while the secondary antibodies were added to *Drosophila* embryos to remove any nonspecific binding. After incubating overnight at 4°C, the oocytes were washed in PTB four times at room temperature. The oocytes were incubated with secondary antibodies for 3-4 hours at room temperature. The oocytes were then washed in PTB and Hoechst33342 (10 μg/mL) was added to stain the DNA. The oocytes were washed twice more in PTB and then were ready for mounting and imaging.

The primary antibodies used in this paper were mouse anti-tubulin monoclonal antibody DM1A at 1:50 conjugated to FITC (Sigma-Aldrich), mouse anti-tubulin monoclonal antibody E7 at 1:200 (Developmental Studies Hybridoma Bank), rat anti-Incenp at 1:600 (Wu et al., 2008), guinea pig anti-CENP-C at 1:1000 (Fellmeth et al., 2023), rat anti-tubulin YOL1/34 at 1:300 (Millipore), rabbit anti-GFP at 1:200 (Thermo Fisher), mouse anti-myc 9E10 at 1:50, Rabbit anti-SPC105R at 1:4000 (Schittenhelm et al., 2009), mouse anti-SPC105R at 1:200 (this study), rat anti-HA (Clone 3F10, Roche) at 1:50, guinea pig anti-MEI-S332 at 1:5000 (Moore et al., 1998), rabbit anti-NDC80 at 1:500, and rabbit anti-WDB at 1:1000 (Sathyanarayanan et al., 2004). The secondary antibodies used were anti-guinea pig 647 at 1:200, anti-rat Cy3 at 1:200, and anti-rabbit 488 at 1:200. All the secondary antibodies were from Jackson ImmunoResearch.

To visualize pairs of homologous chromosomes, fluorescent *in situ* hybridization (FISH) was performed. After fixing the oocytes, 2X SSC was added and the membranes were removed by rolling the oocytes between glass slides. The oocytes were then incubated in increasing concentrations of formamide solution (20%, 40%, and 50%) before being added to the hybridization solution and probes. The probes recognized the 359 bp repeats on the X chromosome (labeled with AlexaFluor 594), the AACAC repeats on the second chromosome (labeled with Cy3), or the dodeca repeats on the third chromosome (labeled with Alexa 647). This was followed by incubation in 80°C for 20 minutes and 37°C overnight. The next day, the oocytes were washed in 50% formamide solution twice at 37°C and 20% formamide once at room temperature. The oocytes were then rinsed three times in 2X SSCT, once in 2X PBST, and incubated in PTB for 4 hours at room temperature. Afterwards, anti-tubulin-FITC antibody was added and the oocytes were incubated overnight at room temperature. The next day, the oocytes were washed in PTB and Hoechst33342 (10 μg/mL) was added. After washing twice more in PTB, the oocytes were ready for mounting and imaging. Oocytes were mounted on a glass slide in SlowFade Gold (Invitrogen) and images were taken with a 63x NA 1.4 lens on a Leica SP8 microscope.

### Imaging Oocytes, Image Analysis, and Statistical Analysis

Images were analyzed using Imaris image analysis software (Bitplane) to count centromere foci, measure karyosome length, KT-MT distances, and intensities. To quantify the distance between centromeres and microtubules, spots marking the centromeres and a surface marking the spindle were created. The Distance Transformation Xtension on Imaris was used to measure the distance outside the surface object. GraphPad Prism software was used to perform all statistical tests, and all experiments were repeated at least twice and analyzed for consistency.

## Acknowledgements

We thank Marina Druzhinina for technical assistance, Christian Lehner and Rager Karess for providing Drosophila stocks and antibodies. Stocks obtained from the Bloomington Drosophila Stock Center (NIH P40OD018537) were used in this study. This work was supported by NIH grant GM101955 to K.S.M.

---

**Figure S 1:**
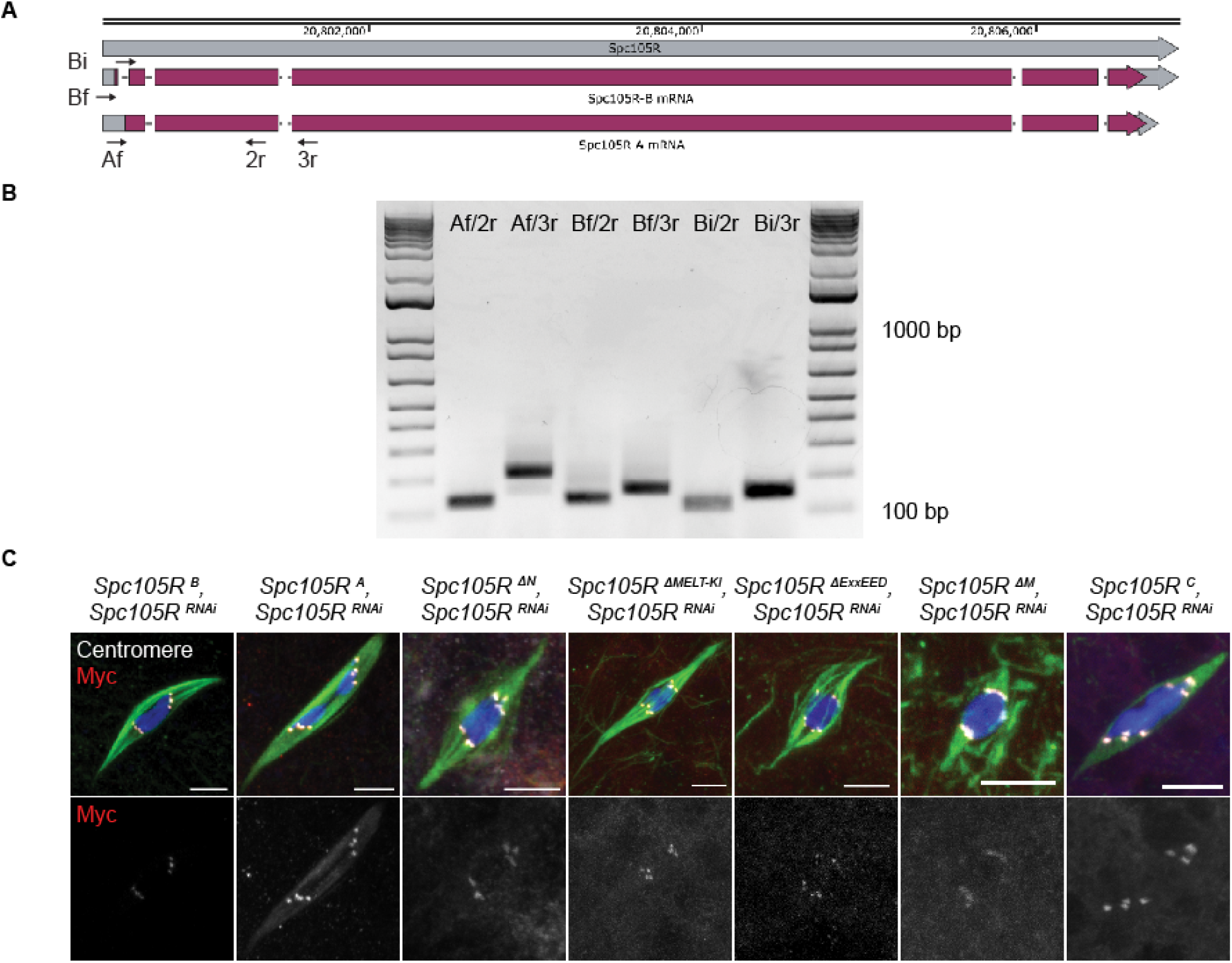
RT-PCR analysis of *Spc105R* isoforms and expression of *Spc105R* deletion mutants. A) Schematic of the two *Spc105R* isoforms showing primers used to amplify in RT-PCR experiments. Primer Bf is in the first exon of the B form but will amplify off both isoforms. Primer Bi spans the intron of the A form and is specific for the B form. Primer Af is within the intron of B form and specific for the A form. Primers 2r and 3r amplify from both splice forms. B) RT-PCR showing amplification with splice-form specific primers. C) Localization of the two SPC105R splice forms and mutants, detected using the MYC epitope (red), with DNA in blue, microtubules in green, CENP-C in white. Scale bar is 5µm.

**Figure S 2:**
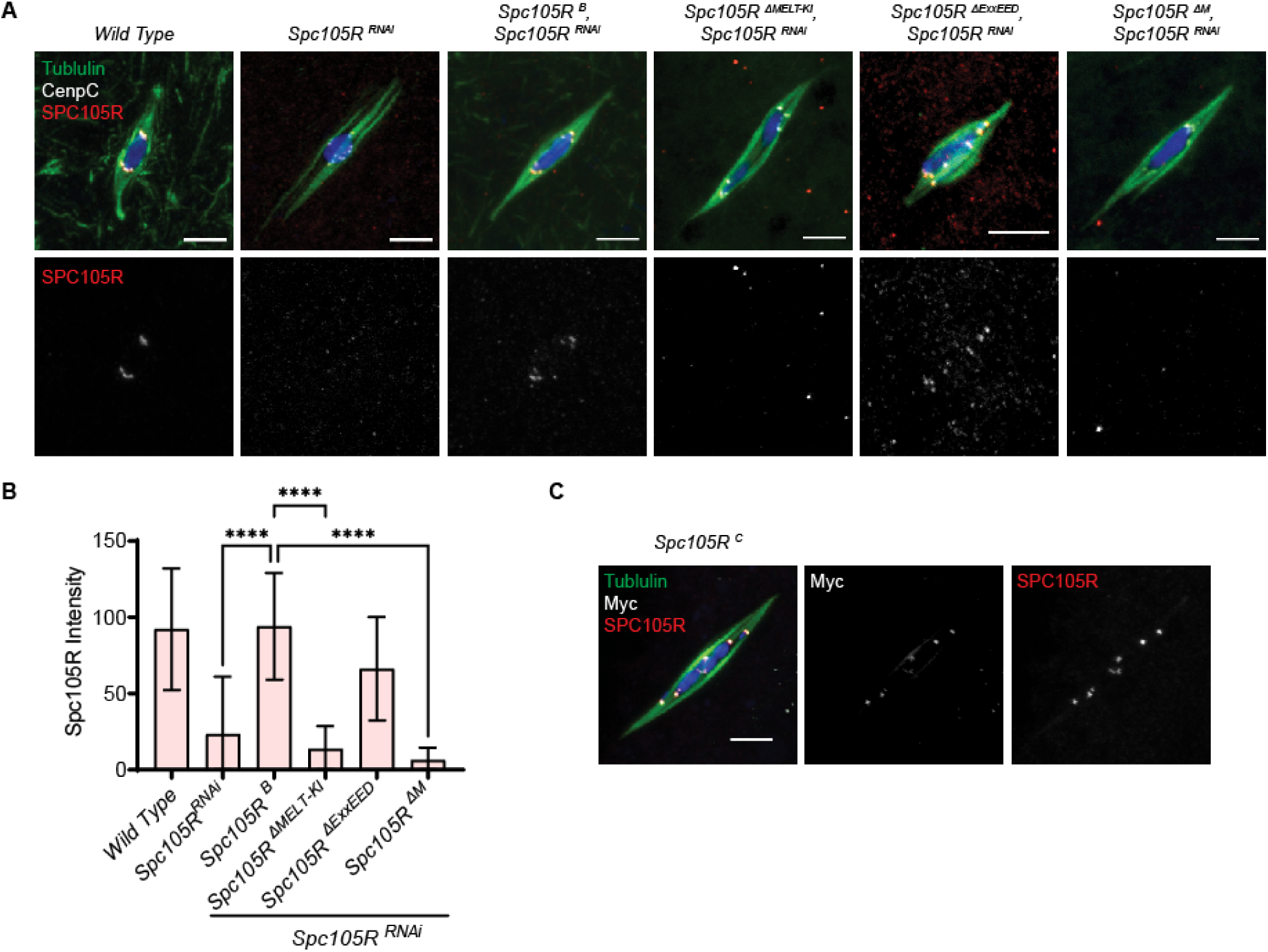
Localization of the SPC105R mutants using the SPC105R antibody. A) SPC105R was detected using an antibody was raised against the first 470 amino acids of the protein (Schittenhelm et al., 2009). DNA in blue, microtubules in green, CENP-C in white, and SPC105R in red. Scale bar is 5 µm. B) Quantification of SPC105R intensity. Error bars indicate standard deviation and a one-sided unpaired t-test was used to show significant decreases in SPC105R intensity (**** = p < 0.0001). Sample sizes are, in order on the graph were n=53, 107, 85, 69, 80, 82. C) *Spc105R^C^* oocytes, with no RNAi and therefore both the transgene (Myc tagged) and endogenous wild-type protein were expressed.

**Figure S 3:**
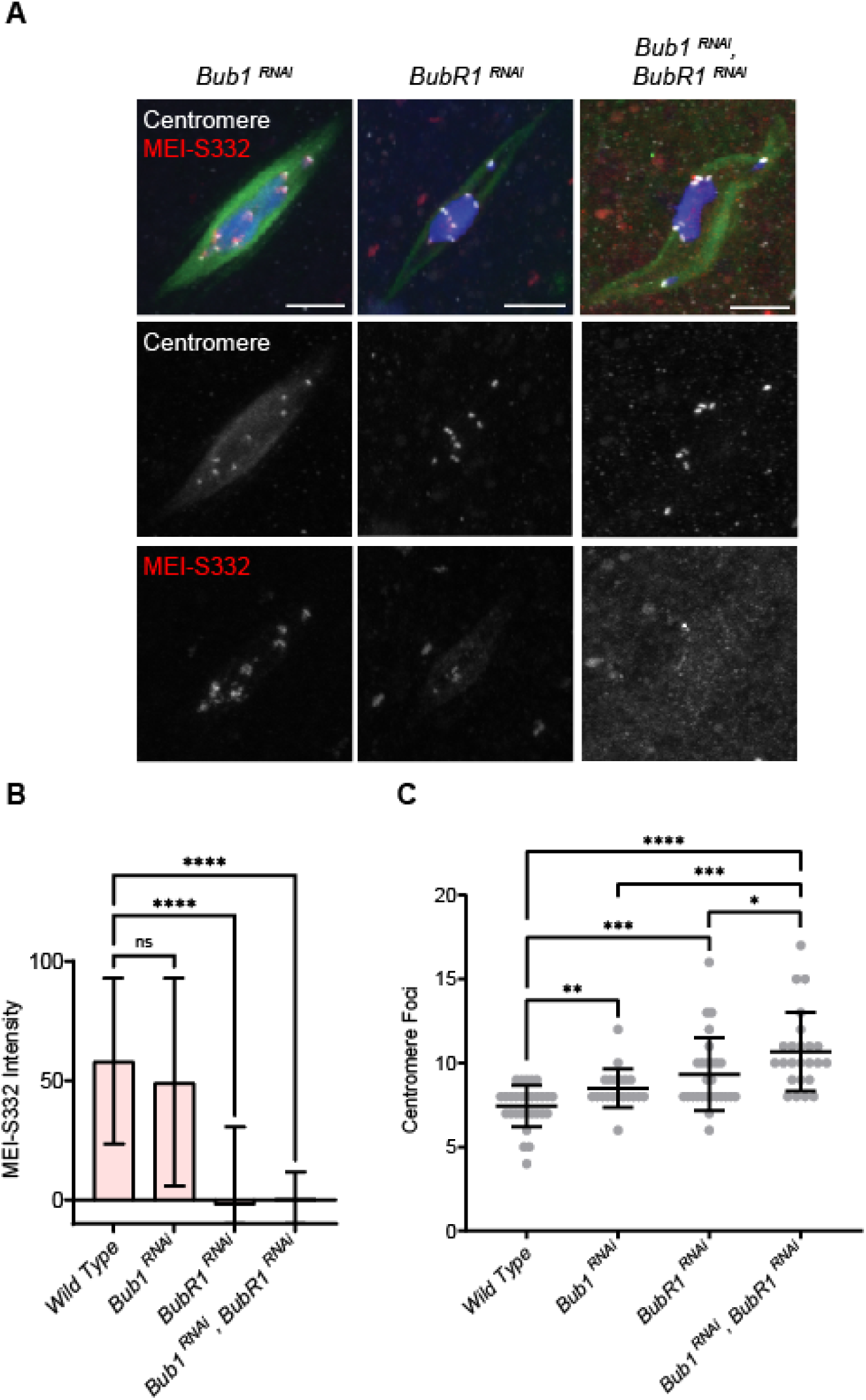
Centromeric cohesion protection in meiosis I. A) Confocal images of Stage 14 *Bub1*^RNAi^, *BubR1*^RNAi^, and *Bub1* ^RNAi^*, BubR1*^RNAi^ oocytes. Merged images show DNA (blue), Tubulin (green), centromeres (white), and the cohesion protection protein MEI-S332 (red). Centromeres and MEI-S332 are shown in separate channels. Scale bars are 5 μm. B) Quantification of MEI-S332 intensity at centromeres, with wild-type (n=71), *Bub1*^RNAi^ (n=104), *BubR1^RNAi^* (n=152) and *Bub1*^RNAi^, *BubR1^RNAi^*oocytes (n=200). C) Quantification of centromere foci, with wild-type (n=29), *Bub1*^RNAi^ (n=20), *BubR1^RNAi^* (n=27) and *Bub1*^RNAi^, *BubR1^RNAi^*oocytes (n=24). Error bars indicate standard deviation, and a ne-sided unpaired t-test was used to show significance, with * = p < 0.02, ** = p = 0.002, *** = p = 0.0002, and **** = p < 0.0001, ns = not significant.

**Figure S 4:**
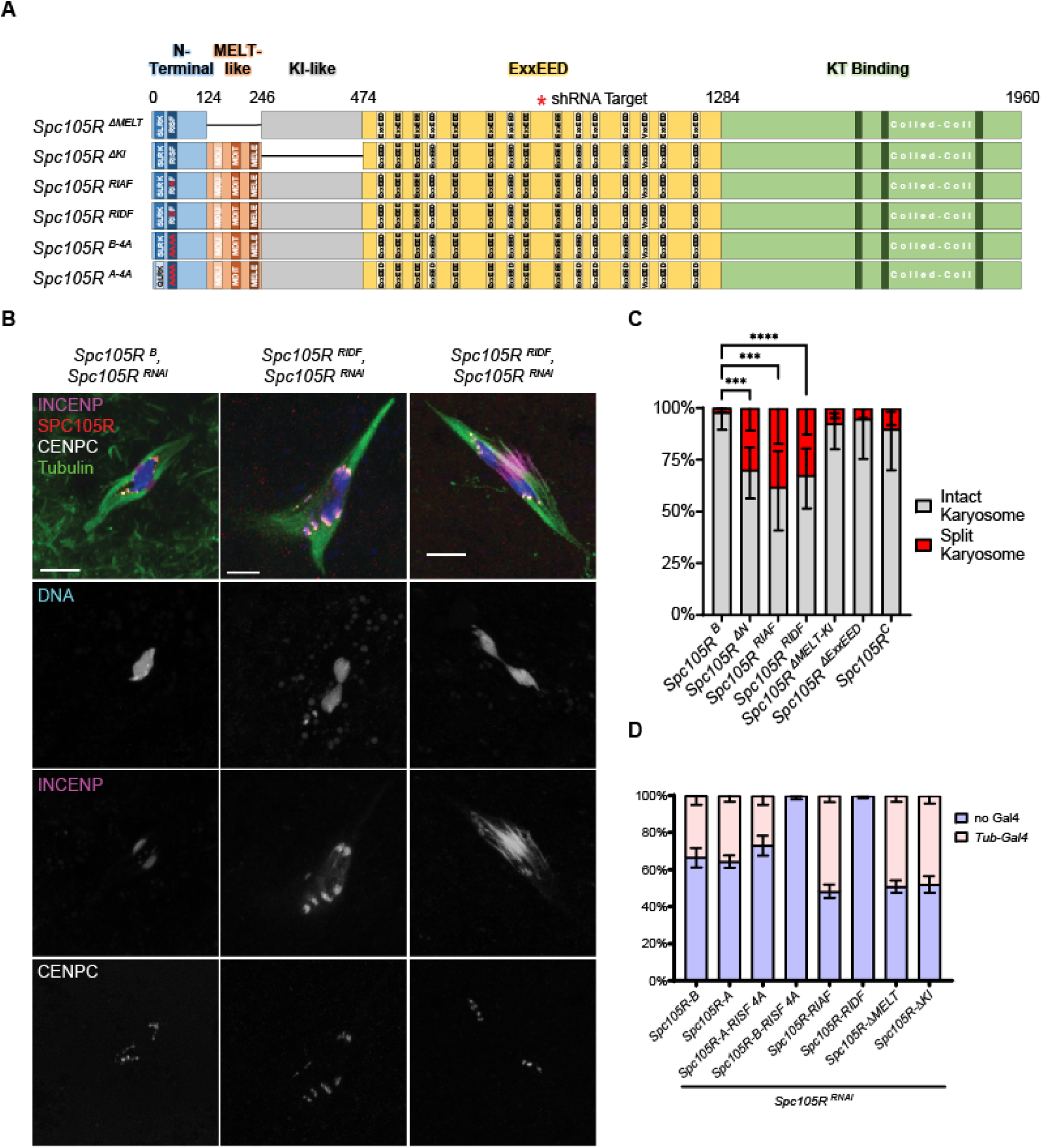
Analysis of RISF, MELT and KI domain mutants A) A schematic of the *Spc105R* mutants effecting the N-terminal domain and the MELT-KI region. The N-terminal includes the SLRK and RISF motifs. B) Confocal images of *Spc105R ^B^* and *Spc105R ^RIDF^* RNAi oocytes. Merged images show DNA (blue), Tubulin (green), CENP-C (white), SPC105R (red), and INCENP (purple). DNA, CENP-C, and INCENP are shown in separate channels. Scale bars are 5 μm. C) Karysome separation phenotype in *Spc105R* mutants. Sample sizes are 50, 40, 20, 50, 19, 21, 37. Fishers exact test when compared to *Spc105R ^B^* : *** = p = 0.001, **** = p < 0.0001. D) Viability of *Spc105R* mutants. Data shows the relative amounts of progeny expressing the *Spc105R^RNAi^*and transgene (*Tub-Gal4)* to siblings that did not (no *Gal4*) (n = >200).

**Figure S 5:**
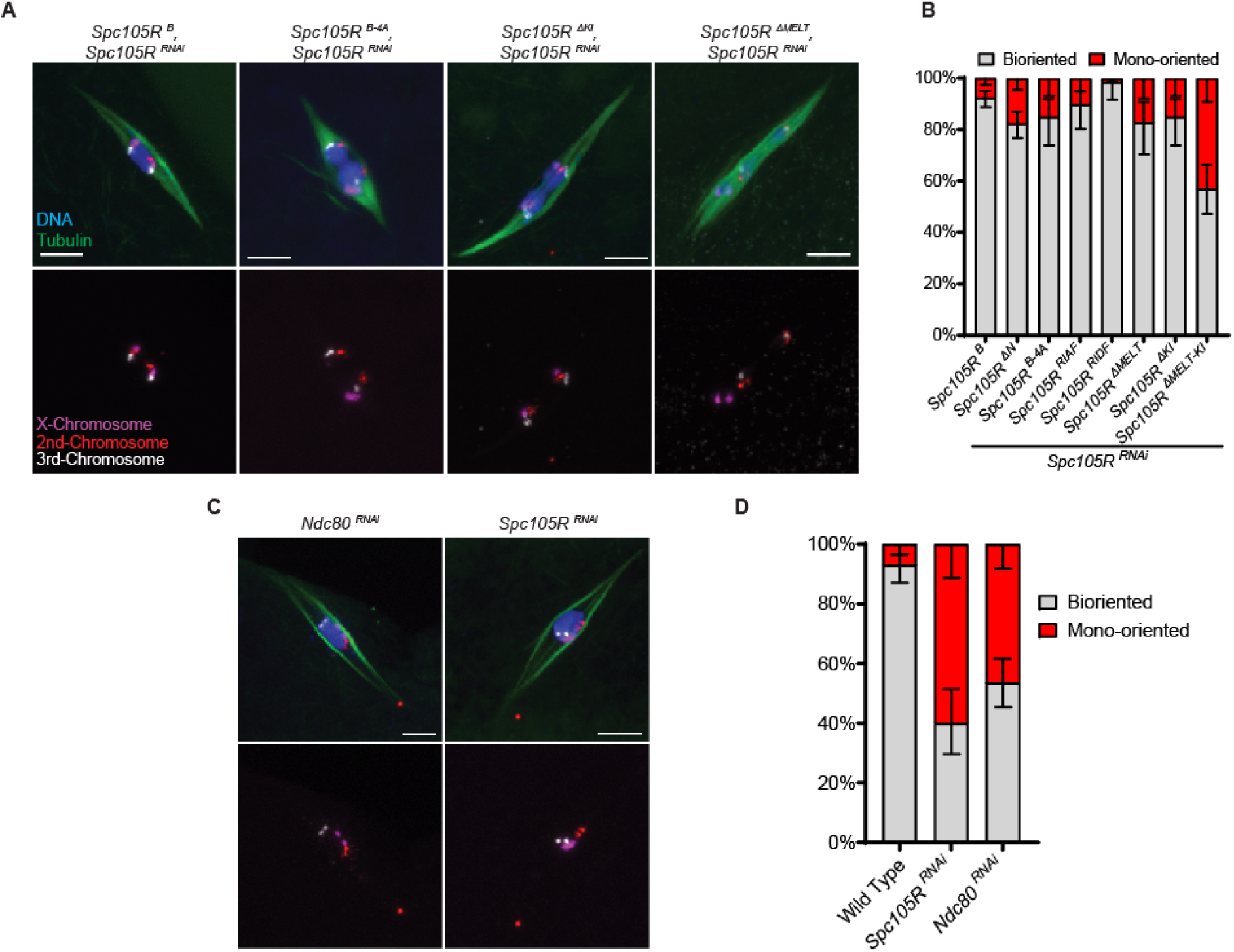
FISH analysis of RISF, MELT, and KI domain mutants A) Confocal images of *Spc105R* RNAi oocytes expressing the indicated transgene. Merged images show DNA (blue), Tubulin (green), the X-Chromosome (purple), the second chromosome (Red), and the third chromosome (white). FISH probes are shown in a separate channel. Scale bars are 5 μm. B) Quantification of percent of chromosome mono-orientation. All mutant transgenes are expressed in *Spc105R^RNAi^*oocytes. Sample sizes are: 275, 204, 60, 68, 63, 52, 60, 100. C) *Ndc80^RNAi^* or *Spc105R^RNAi^*oocytes with the same colors as in A. D) Quantification of percent of chromosome bi-orientation of the oocytes in C. The frequency of mono-orientation was not significantly different between *Ndc80^RNAi^*or *Spc105R^RNAi^* oocytes by Fisher’s exact test (n = 116, 75, 140).

**Figure S 6:**
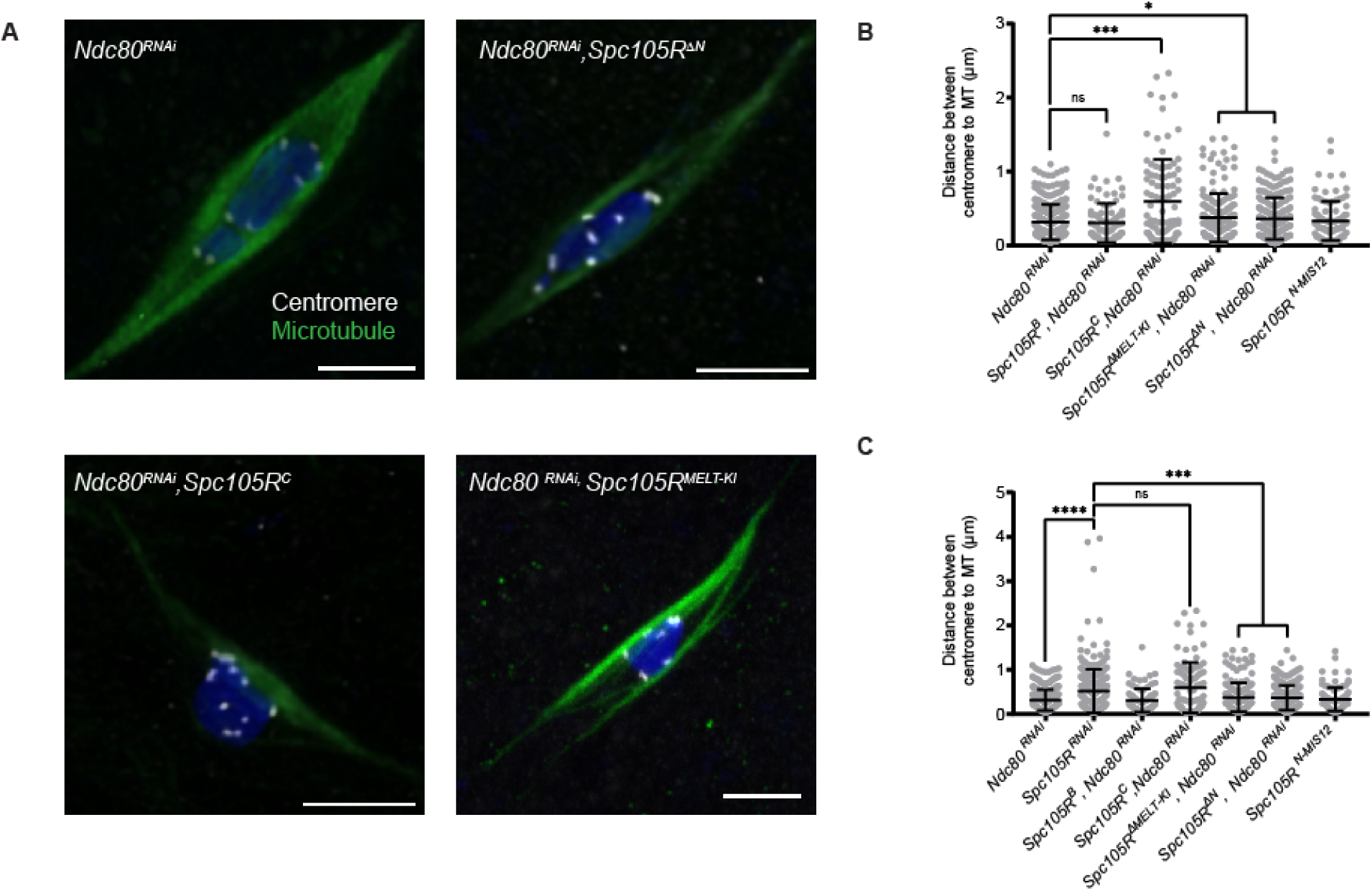
Centromere-microtubule interactions in *Ndc80^RNAi^*, *Spc105R* mutant oocytes (A) Confocal images of Stage 14 *Ndc80^RNAi^*, *Spc105R* mutant oocytes. Merged images show DNA (blue), Tubulin (green), and centromeres (white). Centromeres are shown in a separate channel. Scale bars are 5 μm. B) Distance between centromeres and the nearest microtubules for distances > 0. One-sided unpaired t-test showed significance in number of foci between all channels. The top graph shows comparisons to *Ndc80^RNAi^*(n=325) and the bottom graph shows comparison to *Spc105R^RNA^* (304). Sample sizes for the last 5 on each graph are: 84, 97, 172, 212, 102. Error bars indicate standard deviation, * = p-value < 0.05, ** = p-value < 0.01, **** = p-value < 0.0001, and ns = not significant.

